# Spatio-temporal 3D Mapping of Mouse Cerebellar Vascularization during Embryonic Development

**DOI:** 10.64898/2026.07.16.738922

**Authors:** C Racine, BJ Gonzalez, D Burel

## Abstract

Despite major advances in the study of cerebellar neurogenesis, cerebellar angiogenesis during embryogenesis remains poorly described. Recent advances in tissue clearing, light-sheet microscopy, and artificial intelligence have increasingly enabled detailed 3D modelling of cerebellar vasculature at early developmental stages.

Here, vascular networks in mouse embryos from E11 to birth (P0) were labelled with podocalyxin, αSMA, and PECAM-1 antibodies together with the nuclear marker TO-PRO-3 iodide, cleared, imaged by light-sheet microscopy, and finally modelled and quantitatively analyzed using Imaris and VesselVio.

Our mapping reveals that the three main paired cerebellar arteries—the superior (SCA), anterior inferior (AICA), and posterior inferior (PICA) cerebellar arteries—emerge sequentially between E11 and E13 and display significant topographical variability comparable to that observed in humans. Morphometric analysis demonstrates distinct developmental dynamics, with SCA growth proportional to cerebellar expansion, whereas the AICA and PICA exhibit accelerated extension during later embryonic stages. Interestingly, the PICA does not reach the cerebellum before birth, highlighting the question of its contribution to embryonic cerebellar vascularization. The intrinsic vascular network evolves from a rudimentary bilayer at E11 into a highly branched architecture organized around radial penetrating vessels, giving rise to collaterals that progressively colonized the cerebellar parenchyma during foliation and lobulation. These vascular changes temporally coincided with the successive stages of cerebellar neurogenesis, supporting an interplay between vascular and neuronal development.

Together, our findings provide the first spatio-temporal three-dimensional atlas of cerebellar vascularization during mouse embryogenesis, establishing a reference framework for investigating cerebellar angiogenesis in developmental and pathological conditions.

**Highlights:**

- This work is the first 3D modelling of the cerebellar vasculature in mouse embryo.
- SCA, AICA, PICA develop through distinct spatial and temporal growth programs.
- PICA does not contribute to cerebellar vascularization before birth.
- The intra-cerebellar vascularization evolves at E11 from a simple vessel bilayer.
- Between E13 and P0, radial vessels form collaterals colonizing cerebellum.
- The vascular changes temporally coincided with cerebellar neurogenesis.

## 1. Introduction

The cerebellum is a structure of the central nervous system posterior to the brainstem and anatomically connected to both the pons and the cerebrum. While it has long been considered primarily responsible for motor coordination and control, accumulating evidence now underscores its essential role in higher-order cognitive functions, including learning, language processing, attentional regulation, and emotional processing (Mastrangelo et *al*., 2024; Rondi-Reig et *al*., 2022; Rudolph et *al*., 2023; Van Overwalle, 2024). The adult cerebellum contains nearly 80% of the total neurons in the central nervous system, reflecting this diverse range of functions. Moreover, like the brain, the cerebellum is highly metabolically active and requires a continuous supply of oxygen and glucose to sustain neuronal activity. Thus, a vascular defect occurring in adulthood or during development can lead to significant neurological disorders.

To better understand how vascular alterations may affect cerebellar function, it is first necessary to consider the anatomical structure of the cerebellum. The mature cerebellum is highly organized and can be subdivided into three main regions: the vermis, the cerebellar hemispheres, and the flocculo-nodular lobe. Each of these regions contains three structural components: the cerebellar cortex—characterized by its convoluted surface and subdivision into ten lobules—, the underlying white matter, which contains axonal projections connecting the cortex with the deep cerebellar nuclei and other brain regions, and the deep cerebellar nuclei themselves, which serve as key relay stations for cerebellar output. At a histological level, the cerebellar cortex is also composed of three distinct layers. The innermost layer, called the granular layer, is densely populated with glutamatergic granule cells. These neurons project their axons through the Purkinje cell layer and reach the superficial molecular layer, where they divide to form parallel fibers. The Purkinje cell layer consists of large inhibitory GABAergic neurons, which represent the only output of the cerebellar cortex. These neurons are organized in a single layer and extend their dendritic tree into the molecular layer. Therefore, this latter layer, located just beneath the meninges, contains numerous parallel fibers, the dendritic trees of Purkinje cells, and a variety of GABAergic interneurons (Tamanini et *al*., 2025).

Such a dense and highly specialized cytoarchitecture requires substantial metabolic support throughout the cerebellum. The cerebellar metabolic demand is met through an intricate vascular supply composed of three afferent paired arteries (Naidich et *al*., 2009). The posterior inferior cerebellar arteries (PICA) irrigate the inferior portions of the vermis, the cerebellar hemispheres, and the choroid plexus of the fourth ventricle. The anterior inferior cerebellar arteries (AICA) supply the middle cerebellar peduncle, the flocculo-nodular lobe, the anteroinferior surface of the cerebellum below the horizontal fissure, as well as parts of the inferolateral pons and cranial nerves VII and VIII (Savoiardo et *al*., 1987). Finally, the superior cerebellar arteries (SCA) bifurcate into a medial branch, which irrigates the superior vermis and deep cerebellar nuclei, and a lateral branch, which supplies the superior aspects of the cerebellar hemispheres. Once they reach the surface of the cerebellum, these three vascular afferences divide into several well-defined segments, which ultimately give rise to perforating vessels that penetrate the cortex and irrigate the cerebellum in depth (Blaszczyk et *al*., 2024; Rhoton, 2020).

Although this vascular organization appears relatively stereotyped, many MRI or autopsy studies have revealed a substantial interindividual variability in the origin, branching, distribution, and vascularized territories. The PICA are the most variable, as they can originate from the lateral medullary segment or the premedullary segment of the vertebral arteries (Macchi et *al*., 2004). The AICA are typically described as originating from the basilar artery (BA), but may also emerge from the junction of the basilar and vertebral arteries or from a common trunk with the PICA (Farca-Ureche et *al*., 2005). Finally, although the SCA is considered the most consistent, its morphology can vary, including duplication or triplication originating from the BA (Dağçınar et *al*., 2020). This variability may be attributed to specific mechanisms of vessel selection during embryogenesis, as it has been observed in brain (Lasjaunias et *al*., 2011). However, there is too limited data available regarding the development of the afferent cerebellar arteries to confirm this hypothesis. Recent reports only indicate that the SCA emerges above the rhombencephalon around day 32 of gestation. By day 35, the basilar and vertebral arteries undergo more extensive development, leading to the formation of the PICA and AICA by approximately day 52 (Klostranec and Krings, 2022).

Beyond the variability of the main afferent arteries, even less is known about how the intrinsic vascular network of the cerebellum develops during embryogenesis. One hypothesis suggests that, as in the brain, angiogenesis occurs in parallel of neurogenesis. In the human cerebellum, neuron birth occurs from the 6^th^ week of gestation until the end of the second postnatal year, and involves two primary germinal zones. The ventricular zone gives rise to Purkinje cells and inhibitory interneurons starting from the 6^th^ gestational week, and the rhombic lip generates granule cells around the 8^th^ gestational week (Haldipur et *al*., 2019). These neuronal populations follow distinct migratory pathways. Purkinje cells undergo radial migration to form a monolayer beneath the molecular layer by birth, while granule cells initially migrate tangentially to establish a transient external granular layer, before migrating radially to reach their final position in the granular layer (Leto et *al.*, 2016). Cerebellar neurogenesis is therefore well studied and understood today. In contrast, the mechanisms governing the development of the internal cerebellar vasculature remain poorly characterized in humans and current knowledge is largely extrapolated from cortical and retinal angiogenesis studies, as well as from mouse models. Thus, in mice, the vascular network is described as developing from a superficial perineural vascular plexus, from which vessels invade the neural tissue to provide vascularization. These vessels then contribute to the formation of the internal periventricular plexus, where numerous collateral branches emerge (Karakatsani et *al*., 2019; Vogenstahl,et *al.*, 2022). In the brain, this process occurs between the 4^th^ and 7^th^ weeks of gestation in humans, and between embryonic stages E9 and E13 in mice (Vogenstahl et *al*., 2022). In the retina, it occurs from the 16^th^ week of gestation to the 40^th^ (Selvam et *al*., 2018), and between stages P0 and P21 in mice (Stahl et *al*., 2010).

Altogether, these observations highlight the current lack of knowledge regarding the developmental organization of cerebellar vascularization. Here, we provide a detailed description of cerebellar arterial afferences and the internal vascularization of the cerebellum during embryonic development. Since cerebellar neurogenesis follows the same stages in mice as in humans albeit a shorter frame time (Sepp et *al*., 2024), this description is an essential tool for studying human perinatal vascular accidents involving the cerebellum, such as hemorrhage, infarction (Hortensius et *al*., 2018; Pierson and Al Sufiani, 2016; Villamor-Martinez et *al*., 2019), preeclampsia (Oguz et *al*., 2024) or perinatal hypoxia (Hayakawa et *al*., 2020).

## 2. Materials and methods

### 2.1 Animals

In this study, animals used were wild type C57Bl6/J mouse embryos and pups. The pregnant mice were bred in accredited animal facility (approval number A 76-451-05), in accordance with the French Ministry of Agriculture and the European Community Council Directive 2010/63/UE of September 22^nd^ 2012 relative to protection of animals used for scientific purposes. Mice lived on a 12-hour light/dark cycle, at a constant temperature (21 ± 1°C), with free access to food and water. To collect the embryos and pups at different interest ages, male and females were mixed during 24 hours before isolating females. This isolation day was considered as the 0.5 embryonic day. Mice were daily weight to make sure of the pregnancy. In this study, we chose to focus on the following embryonic ages: the 11^th^, 13^th^, 15^th^ and 17^th^ embryonic days, respectively written E11, E13, E15, E17, and the postnatal day 0 (P0 = birth). The proportion of males and females at each stage was as follows: at E11, 5 males and 4 females; at E13 and E17, 5 males and 5 females; and at E15, 7 males and 5 females. Furthermore, to avoid litter effects, the animals were obtained from different mothers for each age. Depending on the analysis, either all animals or only a subset of them were used. At the interest embryonic day, pregnant mice were sacrificed by decapitation after anesthesia using isoflurane inhalation (Iso-VET 5%). A hysterectomy was performed to collect the uterine horns. The connective membranes covering the embryo and placenta were removed. For stage P0, heads were dissected to extract the entire brain. Embryonic and pup tails were retained for sex determination by *Jarid1* genotyping (Clapcote and Roder, 2005). The samples were rinsed in physiological saline solution (NaCl 9 ‰) before being post-fixed by immersion in 4% paraformaldehyde for 6 hours at 4°C. The embryos were then immersed in phosphate-buffered saline (PBS) at 4°C until use.

### 2.2 Clearing

In this study, the samples were cleared using the adapted iDISCO protocol (Renier et *al*., 2014; 2016). The main steps are the following: 1) Dehydration of the samples by 1h-immersion in increasing concentrations of methanol (MeOH; 20%, 40%, 60%, 80% and 100%), 2) Depigmentation of skin and retina in a solution of 5% hydrogen peroxide (H_2_O_2_) and 95% MeOH, 3) Rehydration by 1h-submersion in decreasing concentrations of MeOH (80%, 60%, 40% and 20%), before rinsing in PTx.2 solution (100 mL PBS 10X, 2 mL Triton-X100s Q.S. 1 L distilled water), 4) Permeabilization in a 1X PBS solution composed of 0.2% TritonX-100, 20% dimethyl sulfoxide (DMSO), glycine (23 mg/mL) and thimerosal at 0.1 g/L, at 37 °C under agitation, 5) Blocking of non-specific binding sites in a 1X PBS solution containing 0.2% Triton-X100, 10% DMSO, 6% Normal Donkey Serum (NDS) and thimerosal for one day at 37 °C under agitation, 6) Incubation of samples with the primary antibodies (Table S1) diluted in PTwH buffer (100 mL PBS 10X, 200 μL Heparin (50 mg/mL), Q.S. 1 L distilled water), with 5% DMSO and 3% NDS at 37 °C under agitation. 7) Six 1-h rinses in the PTwH buffer at room temperature, and 8) Incubation with the appropriate secondary antibodies (Table S1) diluted in PTwH containing 3% NDS at 37°C under agitation. 9) Six new 1-h rinses in PTwH at room temperature under agitation and 10) Dehydration of samples in increasing concentrations of MeOH (20%, 40%, 60%, 80% and 100%). 11) Overnight immersion in a solution containing 66% dichloromethane (DCM) and 33% MeOH to delipidate the tissues and rinse twice for 15 minutes in DCM 100% until the samples dive in the bottom of the tube. 12) Finally, clearing step in dibenzylether (DBE) to homogenize their refractive indexes.

Because of the various sizes of embryos and pups, this protocol was adapted for each age, mainly by optimizing the incubation times and by adding/removing specific steps. Thus, the E13, E15, E17, P0 samples were prepared through the 1^st^ to 10^th^ step. In contrast, to preserve the integrity of E11 embryos, the clearing protocol was started at the 4^th^ step for these samples, and they were embedded in a 1.5% agarose solution between the 9^th^ and 10^th^ step. All these adaptations are available in Table S2.

### 2.3 Imaging / Light Sheet

The 3D acquisitions of the cleared samples were performed using the UltraMicroscope Blaze II (Miltenyi Biotec) using the software Imspector Pro 8.0. The embryos were placed on the holder, lying on their side. They were imaged using one light sheet with horizontal focusing to achieve better spatial resolution across the entire embryo. Each sample was imaged twice: once with a 4× objective lens at zoom 1 to acquire the cerebellum at high magnification, and once with a 4× objective lens at zoom 0.6 to additionally capture the afferent vasculature. Lasers at λ=594 nm and λ=633 nm were used to excite the fluorochromes related to vascular and nuclear markers, respectively. The Z-step between each acquisition was set at 2 µm in depth.

### 2.4 Image processing

OME-Tiff light sheet acquisitions were converted in *.ims* format using Imaris converter (Oxford Instruments) and processed in Imaris software (Oxford Instruments).

#### 2.4.1 Afferent vasculature identification and volume

Using the vascular channel, dedicated regions of interest (ROI) delineating each cerebellar artery were defined. Automated *Surfaces* were then generated based on voxel intensity threshold, enabling the segmentation of those vessels. To obtain vessel length, the *Filament* function was manually applied to trace a segment between the two extremities of each previously generated surfaces. Quantitative measurements were extracted from the software statistics and were then normalized to the cerebellar volumes.

#### 2.4.2 Internal vasculature of the cerebella

To obtain quantitative and comparative vascular parameters, vessel modelling was created using our 3D-image analysis workflow, which we had previously validated (Rodriguez-Duboc et *al*., 2026).

Using Imaris, the cerebellum was manually segmented from the whole hindbrain acquisition using the *Surface* tool by drawing contours every 20 μm in 2D-slice mode, generating a cerebellar mask used for volume calculation. Vascular modelling was then performed on this cerebellar channel using the *Filament* module with the *Automated Autopath Algorithm*, optimized for dense and highly branched vascular networks.

Due to differences in mean vessel diameter and to improve detection accuracy, the vasculature was separated into two subnetworks: superficial and deep networks. Multiscale detection was applied using diameter ranges of 1–10 μm for the deep network and 1–20 μm for the superficial network. This enabled the application of a machine-learning procedure, including orientation change detection, seed point classification, and segment classification, with iterative manual corrections performed until satisfactory modelling was achieved. Once optimized at a given developmental stage on at least three cerebella, the same parameters were applied to all samples of the same stage and subsequently adapted for the other stages. For each network, filament reconstructions were converted into image channels using a Matlab macro linked to Imaris.

To visualize temporal dynamics in the superficial network, automated *Surfaces* reconstructions were generated from the corresponding channel using a constant intensity threshold. The *Morphological Split* function was then applied to automatically separate adjacent or partially connected vessels within the segmented surface by identifying local constriction points and subdividing fused vascular structures into distinct objects to improve vascular segmentation and spatial characterization. A color-coded classification was subsequently performed representing the distance between each segmented surface and the main reference artery, either the SCA or the AICA. To visualize the architecture of the internal vasculature, the resulting filaments were manually color-coded according to their orientation, allowing the identification of distinct vascular organization patterns.

Finally, both networks were exported to Fiji for 8-bit conversion and binarization and saved in *.tiff* and *.nii* formats. The datasets were then analyzed using VesselVio (Bumgarner et *al*., 2021) after verification of skeleton integrity. Quantitative analyses included total vascular volume, surface area, length, number of segments, branchpoints, endpoints, tortuosity, segment partitioning, and mean segment volume, area, length, and radius.

### 2.5 Statistics

Statistical analyses were performed using GraphPad Prism (version 9.0) and R (version 5.1). The normality of arterial volume and length distributions was assessed using the Shapiro–Wilk test. For each analysis, the corresponding statistical approach and its primary objective are briefly described in the table S3. In addition to explore the multivariate structure of the cerebellar vascular network across ages, a principal component analysis (PCA) was conducted using the {FactoMineR} package (Husson et *al*., 2017) on centered and scaled data. Results were visualized using the {factoextra} package (Kassambara et *al*., 2020), with individuals colored according to age. Convex hulls were added to illustrate group dispersion in the principal component space. For all analyses, p-value < 0.05 was considered statistically significant. In figures, asterisks indicate levels of significance: *p-value < 0.05, **p-value < 0.01, and ***p-value < 0.001.

## 3. Results

We developed a 3D imaging approach that enable the modelling and a morphometric analysis of the cerebellar afferent and internal vessels during embryonic development. For each parameter, the sex effect was analyzed. All statistics related to this effect are available in the supplementary data (Table S4).

### 3.1 Topography of the cerebellar afferent arteries during embryonic development

Whole embryo clearing combined with 3D imaging and automated segmentation of the posterior part of the central nervous system enabled the modelling of the cerebellar afferent vasculature. This approach allows a detailed characterization of the topography of the posterior central nervous vascular system at various embryonic stages between E11 to birth in the mouse.

At E11, vascular modelling reveals two prominent, symmetrical paired arteries invading the developing central nervous system from its ventral side. These major vessels correspond to the internal carotid arteries (ICA), followed superiorly by the first extensions of the future posterior communicating arteries in the upper part (PCOMM; Fig. 1). Since these arteries do not directly contribute to cerebellum vascularization, they will not be further described in this article. In contrast, between the two ICA lies the developing basilar artery (BA), which plays a major role in cerebellar blood supply. The caudal part of the BA communicates with ICA via small transient vessels called proatlantal arteries, while its cranial portion is maintained by transitional fetal arteries that are connected to the PCOMM. At this early stage, the BA is not yet well defined but, near the cerebellum, it divides perpendicularly to give rise to the future superior cerebellar arteries (SCA). These arteries develop symmetrically, and nascent vessels arising from these branches organize into a thin vascular leaflet at the surface of the developing cerebellum (Fig. 1).

**Fig. 1.**
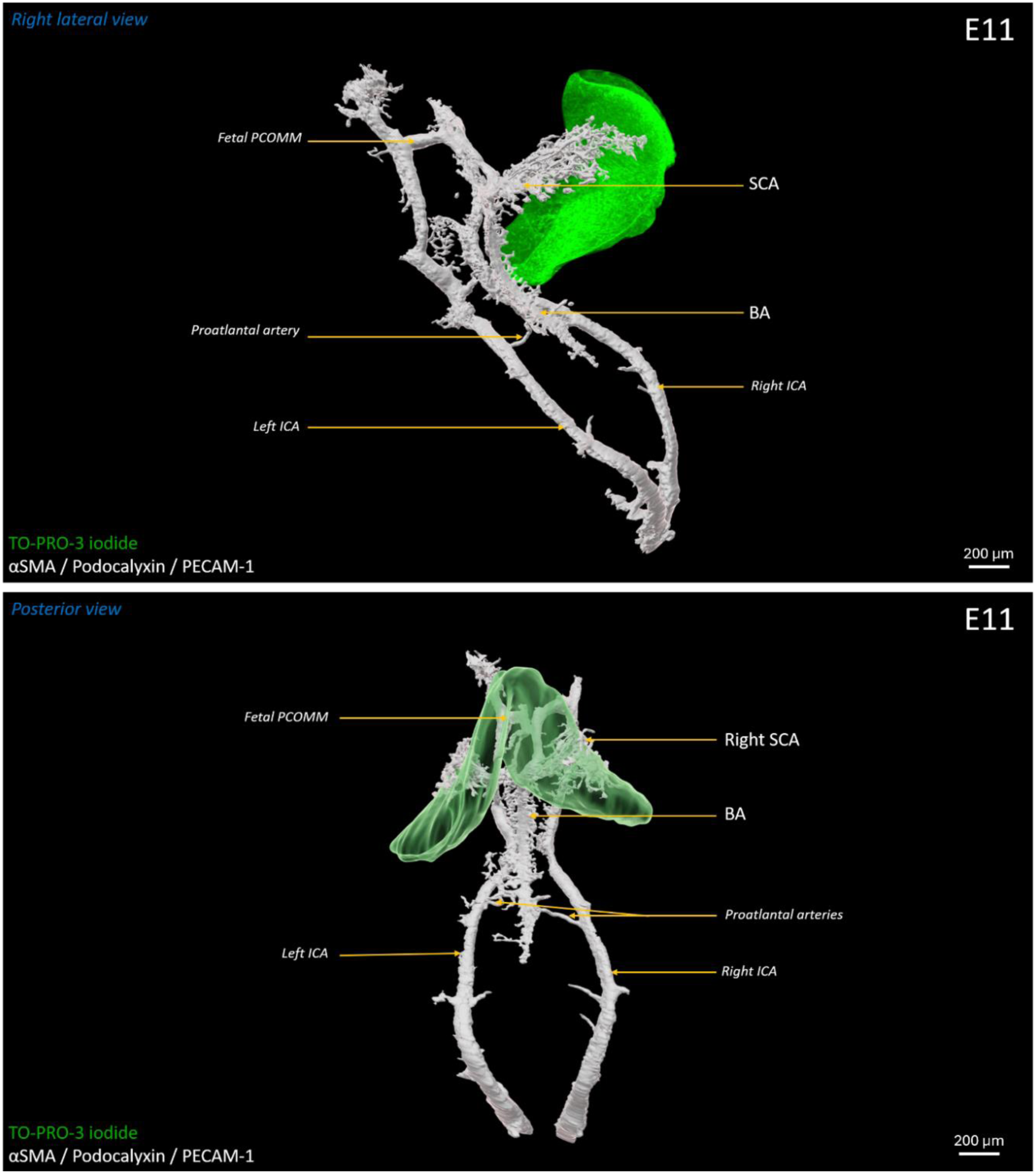
Mapping of the cerebellar afferent arteries in a mouse embryo at E11. 3D modelling of the main arteries involved in cerebellar vascularization in a 11-day-old mouse embryo, shown in right lateral view (top) and posterior view (bottom). Rhombomere 1, corresponding to the prospective cerebellum, was labeled with the DNA intercalating dye TO-PRO-3 iodide and is shown in green. Blood vessels were identified by triple immunolabeling for αSMA, podocalyxin, and PECAM-1 and are shown in white. Arterial labels shown in larger font correspond to vessels directly contributing to cerebellar blood supply, whereas labels shown in smaller italicized font indicate arteries not directly involved in cerebellar vascularization. αSMA: alpha-smooth muscle actin; BA: basilar artery; ICA: internal carotid artery; PCOMM: posterior communicating artery; PECAM-1: platelet endothelial cell adhesion molecule; SCA: superior cerebellar artery.

At E13, a clear separation emerges between the ICA–PCOMM system and the BA, associated with the regression of the proatlantal arteries and the disappearance of the transient fetal arteries between PCOMM (Fig. 2). As a result, the BA becomes a distinct and well-defined structure. Its caudal part undergoes substantially growth and divides to form the vertebral arteries (VA). The primitive posterior inferior cerebellar arteries (PICA) emerge from the posterior branches of these VA, while several vascular anlages sprout from the BA. Notably, in the middle of the BA, two curved paired arteries appear, corresponding to the primitive anterior inferior cerebellar arteries (AICA). In parallel, at the cranial part of the BA, the primary SCA initially follow a straight course before diverging laterally and medially in two branches that extend dorsally toward the posterior cerebellum. Concurrently, the previously flat vascular structures observed at E11 in the distal ends of the SCA become less prominent. Additionally, at the distal part of the BA, the posterior cerebral arteries (PCA) begin to emerge (Fig. 2).

**Fig. 2.**
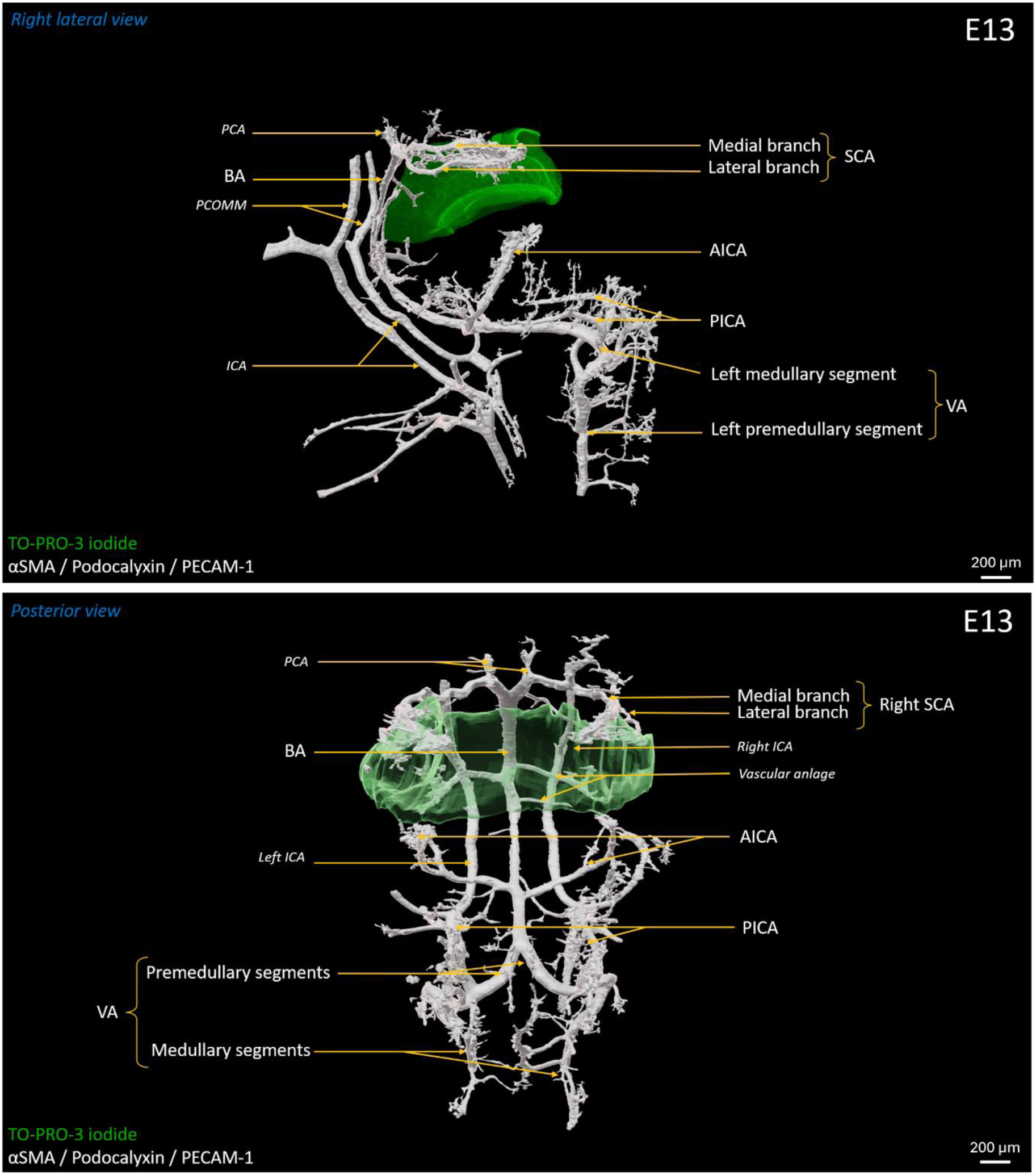
Mapping of the cerebellar afferent arteries in a mouse embryo at E13. 3D modelling of the main arteries involved in cerebellar vascularization in a 13-day-old mouse embryo, shown in right lateral view (top) and posterior view (bottom). The cerebellum was labeled with the DNA intercalating dye TO-PRO-3 iodide and is shown in green. Blood vessels were identified by triple immunolabeling for αSMA, podocalyxin, and PECAM-1 and are shown in white. Arterial labels shown in larger font correspond to vessels directly contributing to cerebellar blood supply, whereas labels shown in smaller italicized font indicate arteries not directly involved in cerebellar vascularization. AICA: anterior inferior cerebellar artery; αSMA: alpha-smooth muscle actin; BA: basilar artery; PCA: posterior cerebral artery; PCOMM: posterior communicating artery; PECAM-1: platelet endothelial cell adhesion molecule; PICA: posterior inferior cerebellar artery; ICA: internal carotid artery; SCA: superior cerebellar artery; VA: vertebral artery.

At E15, the vascular network exhibits a more organized and clearly defined architecture. The PICA show marked growth and extend bilaterally along a course parallel to the BA (Fig. 3). The AICA retain their curved trajectory and reach the ventral part of the cerebellum, although they do not yet establish clear connections. Symmetrical transverse vascular sprouts are still present along the BA, but they remain poorly developed. Meanwhile, the branches of the SCA continue to expand both laterally and medially into the cerebellar parenchyma, progressively recovering the developing cerebellum (Fig. 3).

**Fig. 3.**
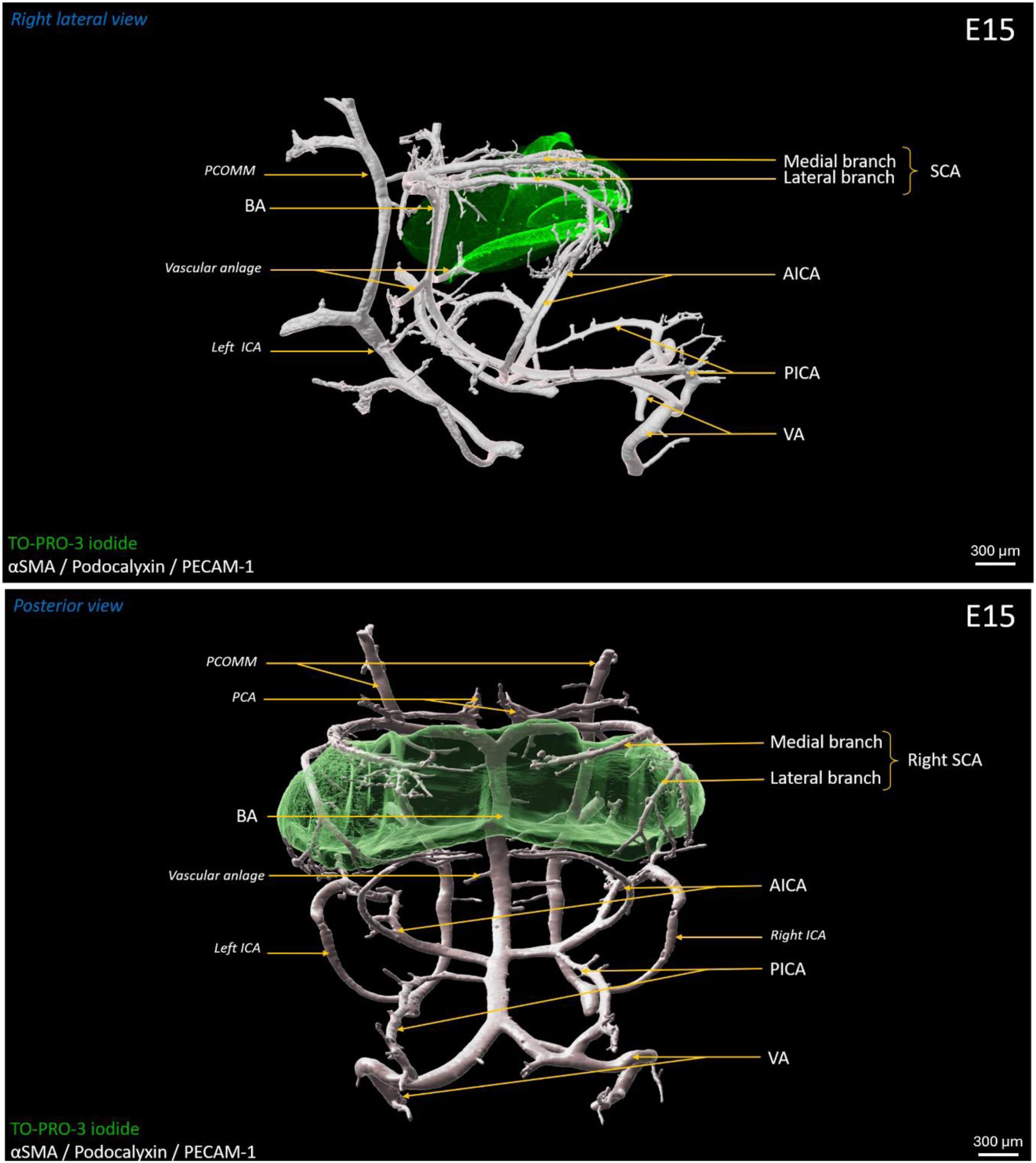
Mapping of the cerebellar afferent arteries in a mouse embryo at E15. 3D modelling of the main arteries involved in cerebellar vascularization in a 15-day-old mouse embryo, shown in right lateral view (top) and posterior view (bottom). The cerebellum was labeled with the DNA intercalating dye TO-PRO-3 iodide and is shown in green. Blood vessels were identified by triple immunolabeling for αSMA, podocalyxin, and PECAM-1 and are shown in white. Arterial labels shown in larger font correspond to vessels directly contributing to cerebellar blood supply, whereas labels shown in smaller italicized font indicate arteries not directly involved in cerebellar vascularization. AICA: anterior inferior cerebellar artery; αSMA: alpha-smooth muscle actin; BA: basilar artery; PCA: posterior cerebral artery; PCOMM: posterior communicating artery; PECAM-1: platelet endothelial cell adhesion molecule; PICA: posterior inferior cerebellar artery; ICA: internal carotid artery; SCA: superior cerebellar artery; VA: vertebral artery.

At E17, vascular remodeling continues, leading to the emergence of more robust and well-defined arterial structures. The PICA maintain their course parallel to the BA (Fig. 4). They do not yet extend toward the developing cerebellum but remain oriented toward the AICA. No marked changes are observed in the SCA compared with E15, except that the vessel density within the vascular leaflet appears reduced. Overall, the vascular architecture continues to mature while preserving the general organization established at the previous stage.

**Fig. 4.**
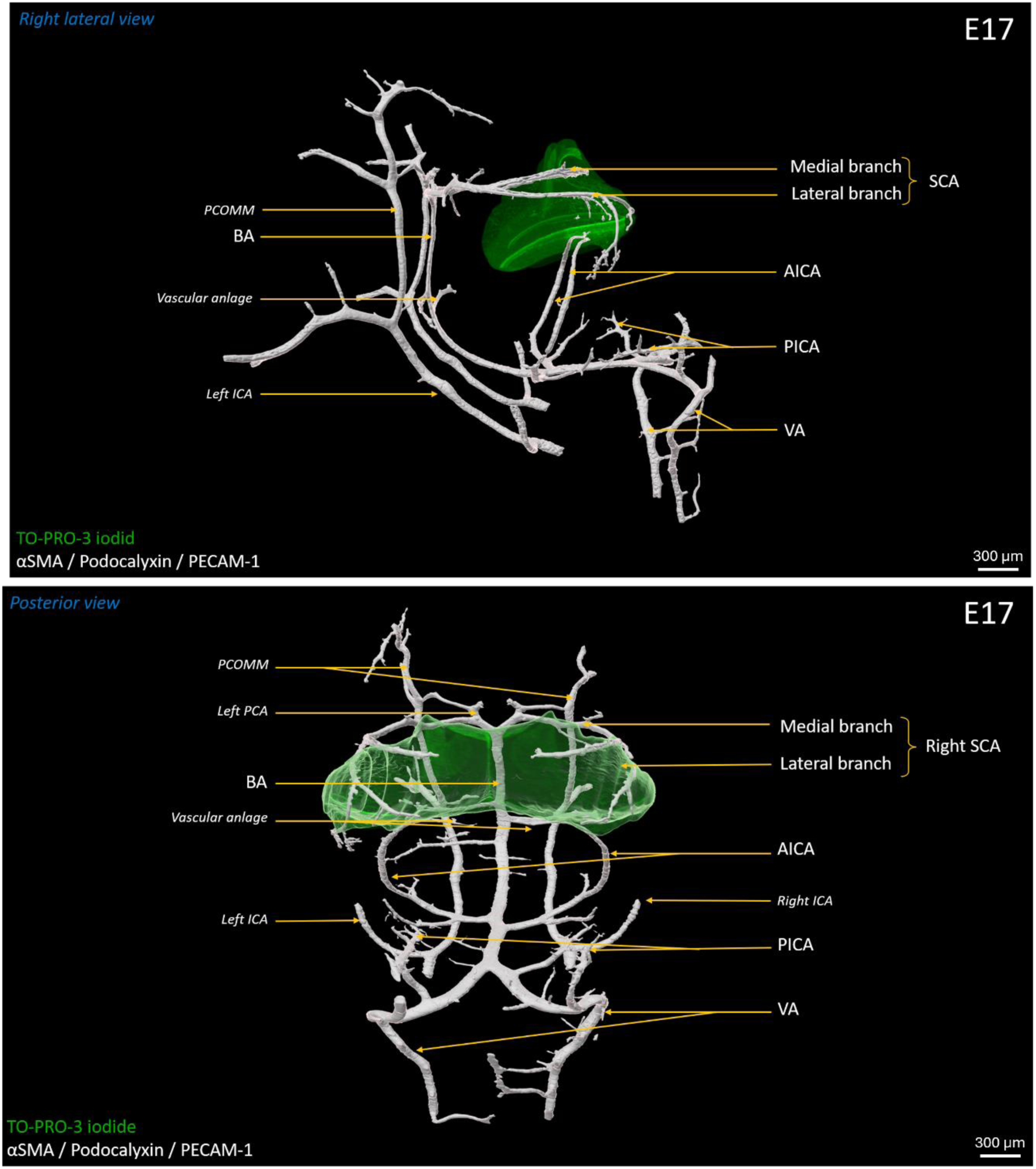
Mapping of the cerebellar afferent arteries in a mouse embryo at E17. 3D modelling of the main arteries involved in cerebellar vascularization in a 17-day-old mouse embryo, shown in right lateral view (top) and posterior view (bottom). The cerebellum was labeled with the DNA intercalating dye TO-PRO-3 iodide and is shown in green. Blood vessels were identified by triple immunolabeling for αSMA, podocalyxin, and PECAM-1 and are shown in white. Arterial labels shown in larger font correspond to vessels directly contributing to cerebellar blood supply, whereas labels shown in smaller italicized font indicate arteries not directly involved in cerebellar vascularization. AICA: anterior inferior cerebellar artery; αSMA: alpha-smooth muscle actin; BA: basilar artery; PCA: posterior cerebral artery; PCOMM: posterior communicating artery; PECAM-1: platelet endothelial cell adhesion molecule; PICA: posterior inferior cerebellar artery; ICA: internal carotid artery; SCA: superior cerebellar artery; VA: vertebral artery.

At birth, numerous collaterals arising from the medial and lateral branches of the SCA encircle the cerebellum (Fig. 5). The AICA form well-developed connections with the anteroinferior region of the cerebellum, reflecting its advanced maturation. The course of the PICA is more variable. As illustrated in Fig. 5 (posterior view), some PICA establish a direct fusion with the BA at the level of the AICA origin, whereas others remain independent and give rise to numerous collateral branches that either anastomose with the AICA or extend dorsally toward the hindbrain.

**Fig. 5.**
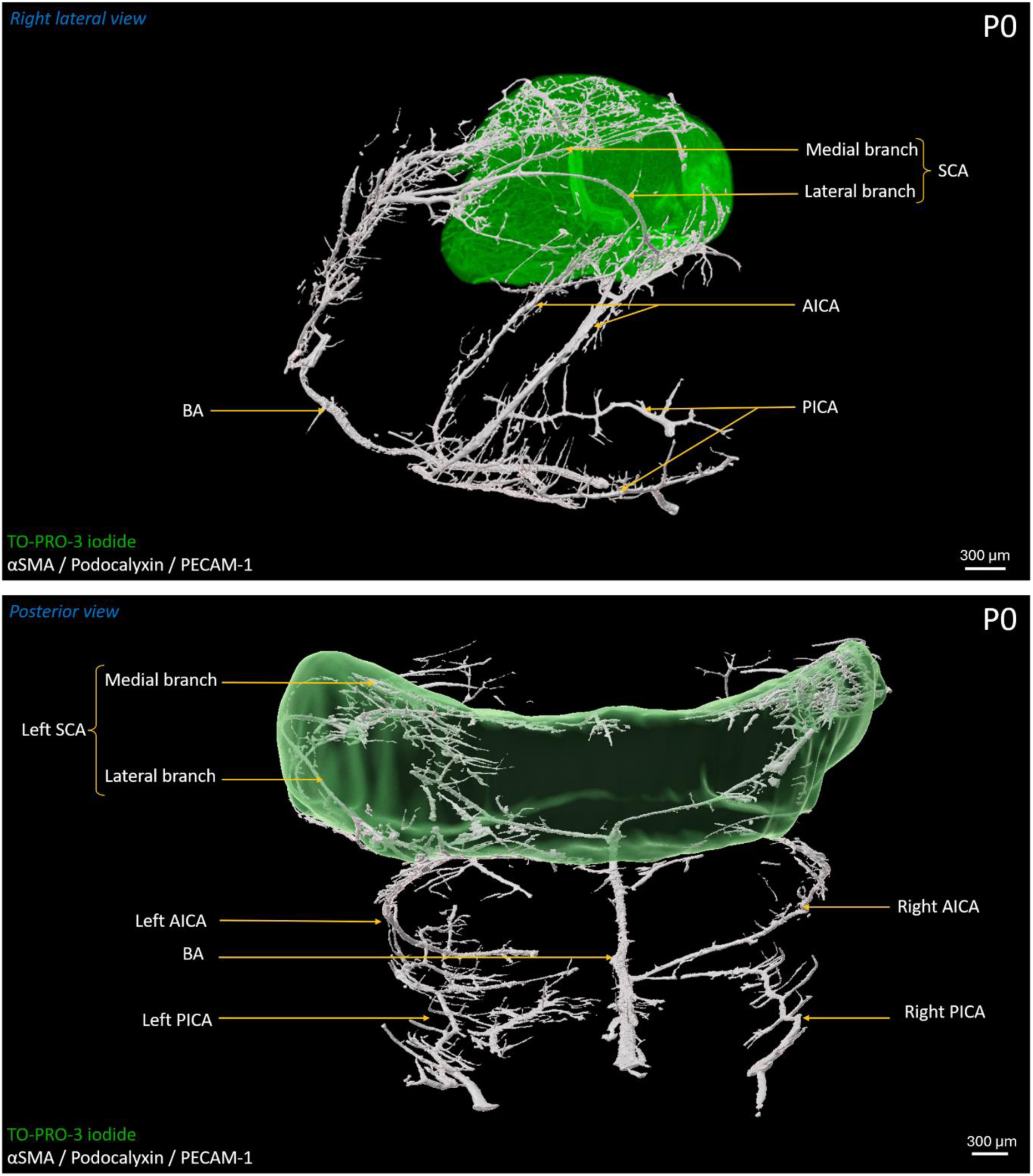
Mapping of the cerebellar afferent arteries in a newborn (P0) mouse. 3D modelling of the main arteries involved in cerebellar vascularization in a newborn mouse, shown in right lateral view (top) and posterior view (bottom). The cerebellum was labeled with the DNA intercalating dye TO-PRO-3 iodide and is shown in green. Blood vessels were identified by triple immunolabeling for αSMA, podocalyxin, and PECAM-1 and are shown in white. Arterial labels shown in larger font correspond to vessels directly contributing to cerebellar blood supply, whereas labels shown in smaller italicized font indicate arteries not directly involved in cerebellar vascularization. AICA: anterior inferior cerebellar artery; αSMA: alpha-smooth muscle actin; BA: basilar artery; PECAM-1: platelet endothelial cell adhesion molecule; PICA: posterior inferior cerebellar artery; SCA: superior cerebellar artery.

### 3.2 Morphometric analysis of the afferent cerebellar arteries during embryonic development

Following the mapping of cerebellar arteries throughout embryonic development, a quantitative analysis was performed to evaluate their morphometric evolution. The use of 3D vascular modelling with Imaris enabled the extraction of structural parameters from automatic surfaces. Thus, for each embryonic stage studied (namely E11, E13, E15 and E17), arterial volume and length were measured and normalized to cerebellar volume, allowing the analysis of growth dynamics across time.

First, the segmented arterial models were distinctly color-coded to facilitate identification and improve visual interpretation across samples (Fig. 6A). The automatic arterial segmentations were then analyzed to extract volume and length data for each artery. The ratio of the total arterial volume to cerebellar volume showed a significant increase between E11 and E13 (*p-value = 0.0248) and between E11 and E15 (***p-value = 0.0008; Fig. 6B). Moreover, at E11, the total SCA volume of the SCA appears higher in male than in female, mainly due to the left SCA (Table S4). In contrast, no significant difference was observed between stages from E13, indicating a balanced growth between the arterial network and the cerebellum (Table S5).

**Fig. 6.**
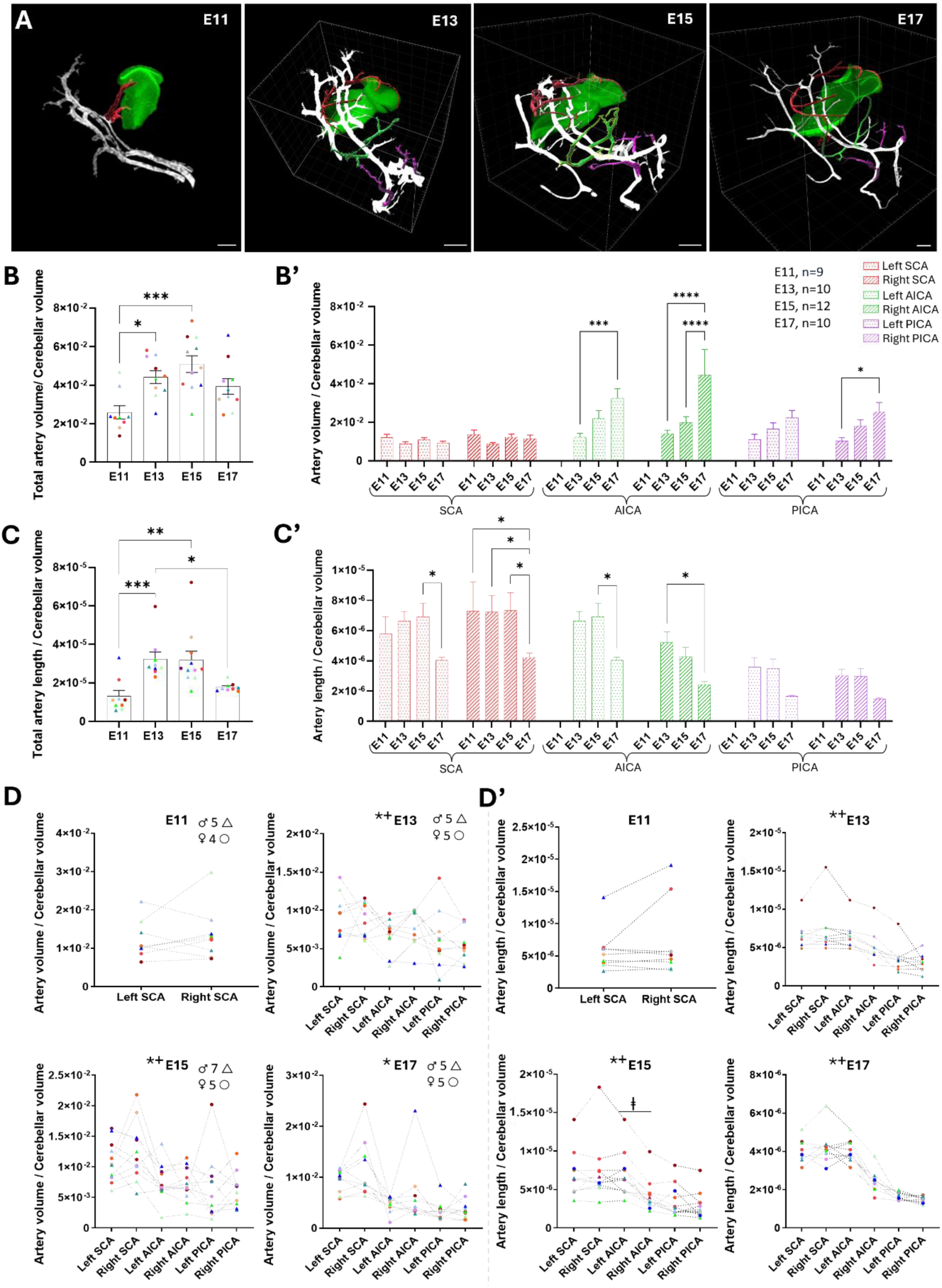
Evolution of the morphology of the three paired cerebellar arteries during mouse embryonic development. (A) 3D modellings illustrating the mapping of the SCA (in red), AICA (in green) and PICA (in purple) at E11, E13, E15 and E17. Scale bars: 200 µm. (B, B’) Histograms showing the total volume of three cerebellar arteries (B) and the average volume of each cerebellar artery (B’) relative to the volume of the cerebellum at E11, E13, E15 and E17. (C, C’) Histograms indicating the total length of three cerebellar arteries (C) and the average length of each cerebellar artery (C’) relative to the volume of the cerebellum at E11, E13, E15 and E17. Data are expressed as mean +/- SEM. *p-value < 0,005; ***p-value < 0,001; ****p-value < 0,0001. (D, D’) Charts detailing the total volume (D) and the total length (D’) of the three paired cerebellar arteries per individual in E11, E13, E15 and E17 mouse embryos. Color indicates the sample identity and triangles represent males and circles represent females. Asterisks indicate statistically significant inter-artery differences, crosses indicate significative inter-individual differences and double cross indicates significative difference between left and right artery. AICA: anterior inferior cerebellar artery; PICA: posterior inferior artery; SCA: superior cerebellar artery.

To determine whether all arteries exhibit a similar growth pattern during embryonic development, a more detailed analysis was performed by quantifying the volume of each cerebellar artery individually, with a distinction between the right and left branches. Over time, the volume of the SCA showed no variability depending on cerebellar volume, indicating that arterial volume increases proportionally with cerebellum growth (Fig. 6B’). In contrast, the AICA exhibited significant continuous volumetric growth during development, with increases of 62% in the left branch (***p-value = 0.0006) and 68% in the right branch (****p-value < 0.0001) between E13 and E17. Similarly, the PICA showed continuous growth between E13 and E17, with a more pronounced and statistically significant increase for the right branch (+60%; *p-value = 0.0175). Thus, the growth rate of AICA and PICA is higher than that of the cerebellum during embryogenesis.

Concerning artery length, which represents a reliable indicator of the dynamics of cerebellar vascularization, the ratio of total arterial length to cerebellar volume showed a significant increase between E11 and E13 (***p-value = 0.0007), as well as between E11 and E15 (**p-value = 0.0037; Fig. 6C). This increase stabilized between E13 and E15 before decreasing at E17 (*p-value = 0.0185; Fig. 6C). Interestingly, taken together, the three pairs of arteries have a greater total length in females than in males at E15, due to the left-sided component of these vessels (Table S4). As for arterial volume, a detailed characterization of the arteries was performed. It reveals that both the left and right SCA exhibited a significant 42% decrease in their length relative to cerebellar volume between E15 and E17 (left: *p-value = 0.0265; right: *p-value = 0.0141; Fig. 6C’). For the AICA, a progressive decrease in the ratio between arterial size and cerebellar volume was observed for both the right and left branches between E13 and E17. This reduction is statistically significant for the right AICA (−53%, *p-value = 0.0408), while it does not reach significance for the left AICA (−39%, p-value = 0.0651 - ns). A similar trend was noted for the PICA, but the changes do not reach statistical significance (Table S5).

Finally, to evaluate the variability of cerebellar artery development, arterial volume and length measurements were plotted for each individual sample (Fig. 6D, D’). This representation enabled the assessment of both inter-artery and inter-individual variability through a mixed-effects analysis, in which artery was considered as the treatment factor and individual embryos as the matching factor (Table S5). Our results show that the volume and length of right and left SCA did not significantly differ at E11. In contrast, from E13 to E17, a significant overall difference among arteries was observed for both volume and length (mixed-effects analysis; treatment; 0.0006 < p-value < 0.0172; Fig. 6D, D’). More specifically, at E15, the left AICA appeared significantly larger than the right AICA (Friedman test with Dunn’s post hoc test; *p-value = 0.0157; Fig. 6D’). During the same developmental period, significant inter-individual variability was also detected for both parameters, except for arterial volume at E17 (mixed-effects analysis; matching effect; 0.0001 < p-value < 0.0069; Fig. 6D’). Moreover, analysis of inter-individual and inter-arterial variance components indicates that, for volume, variability between arteries was the main source of variation. In contrast, for length, inter-individual variability represented the primary contributor to the observed dispersion (Table S5).

### 3.3 Topographic variations of mouse cerebellar arteries

The wide topographic variability and branching patterns of the cerebellar arteries described in humans (Błaszczyk et *al*., 2024) led us to further extend our study with a qualitative analysis of cerebellar vascularization, with a particular focus on the AICA and PICA since the SCA are known to exhibit a more homogeneous pattern.

The typical arterial topography in humans is characterized by single AICA and PICA arising from the BA and the VA, respectively (Fig. 7A). However, many other configurations are frequently observed such strictly duplicated vessels, duplicated-fused vessels, meaning that two primordial branches subsequently merge, and branching between arteries before continuing their course towards the cerebellum (Fig. 7A). A similar variability was observed in mouse embryos. As an illustration, the provided modelling of the cerebellar afferent vascularization in an E15 embryo shows a duplicated and fused left PICA (Fig. 7B^3^), a branching between the left PICA and the BA (Fig. 7B^5^), a duplicated and fused right AICA (Fig. 7B^2^), and a branching between the right PICA and the right AICA ((Fig. 7B^4^).

**Fig. 7.**
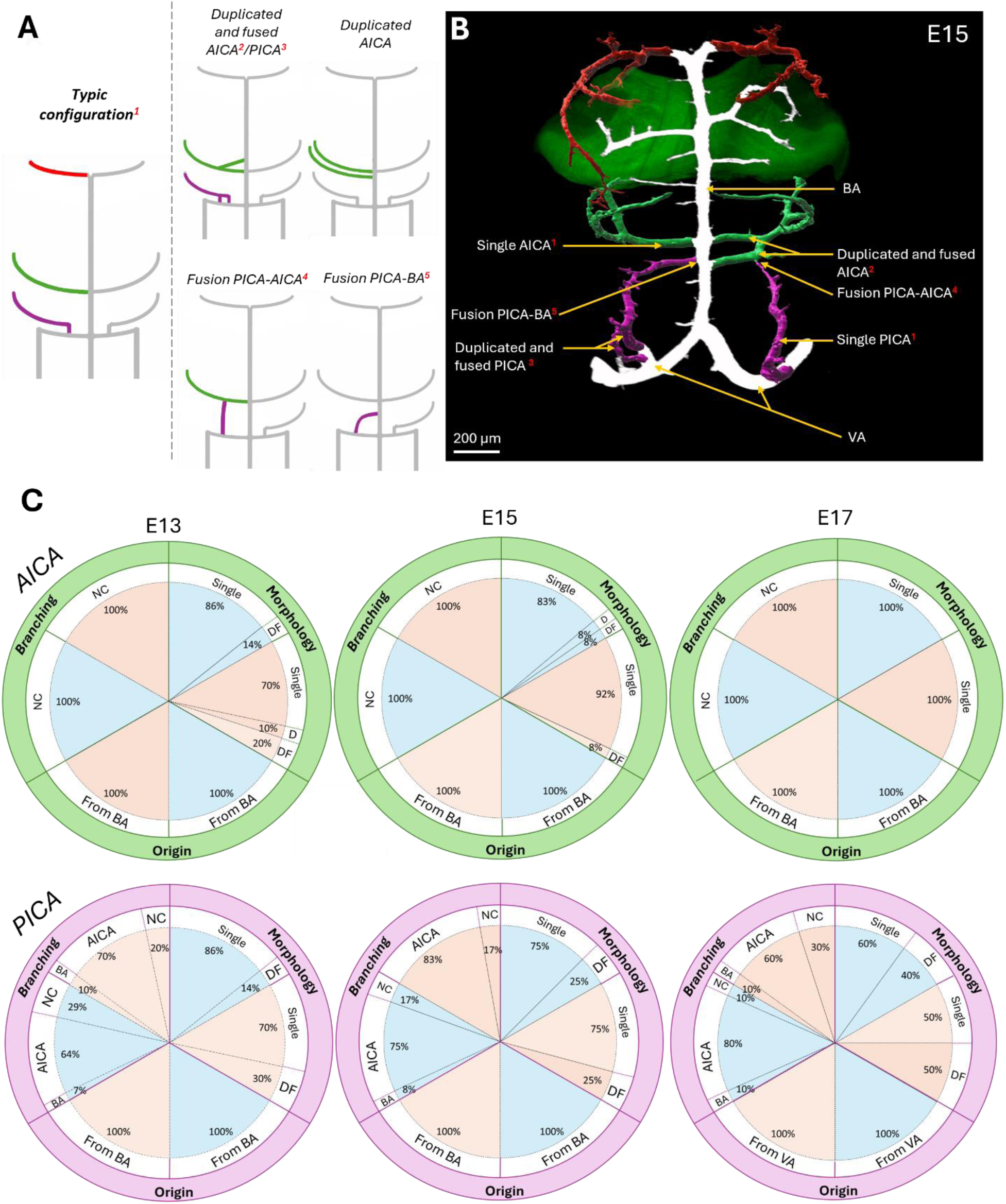
Analysis of the topographic variability of the three cerebellar arterial pairs. (A) Schematic representation of the typic configuration of cerebellar arterial topology and the main variations in morphology and branching observed in 30 embryos at different developmental stages. (B) Illustration of an atypical configuration of the cerebellar arteries observed in a mouse embryo at E15. The PICA are shown in purple, the AICA in green, and the SCA in red. This 3D modelling of the main cerebellar arteries clearly shows that the left PICA duplicated before refusing (3), then fused with the basilar artery (BA; 5), that the right PICA fused with the right AICA (4), and that the right AICA duplicated and refused before reaching the cerebellum (2). (C) Pie charts illustrating the distribution of PICA and AICA variants observed in the mouse embryos at E13 (n=12; ♂=7; ♀=5), E15 (n=10; ♂=5; ♀=5) and E17 (n=10; ♂=5; ♀=5), based on their origin, morphology (single or duplicated and fused) and branching pattern (normal course, to the BA or to the AICA). For each variant, the percentage observed in male (blue) and female (orange) are detailed. AICA: anterior inferior cerebellar artery; BA: basilar artery; D: duplicated; DF: duplicated and fused; NC: normal course; PICA: posterior inferior artery; SCA: superior cerebellar artery; VA: vertebral artery.

The complete inventory of morphological and branching variations was compiled across all our samples, with female and male embryos analyzed separately (Fig. 7C). Across all samples examined, the AICA consistently originated from the BA. Their morphology exhibited a stage-dependent pattern, with a higher variability at E13 that tended to decrease at E17 (Fisher’s exact test; p-value =0.0534 - ns). At E13, 14% of AICA in males displayed a duplicated-fused pattern, whereas females exhibit 30% of either strictly duplicated or duplicated-fused arteries. In contrast, at E17, all AICA appeared as single branches. No significant sex-related differences were detected in AICA morphological variability (Table S4).

For the PICA, their origin from the VA remained consistent throughout development. Two morphological patterns, single and duplicated-fused, were observed at each stage. Regarding branching configuration, connections frequently occurred with the AICA and less frequently with BA, with these patterns remaining relatively stable over time. No significant sex-related effects were detected for any of the topographic parameters of the PICA (Table S4).

### 3.4 Macro-architecture of the vasculature of the developing cerebellum

#### 3.4.1 Distribution of the vascular contribution of the main arteries

To characterize cerebellar vascular development, both superficial and internal vascular networks were identified and analyzed separately. First, a distance-based color map was applied to visualize the contribution of the major arteries to the formation and expansion of the superficial vascular network during cerebellar development (Fig. 8). Based on the maturation pattern of the afferent vasculature described above (Fig. 1-5), only the contributions of the SCA and the AICA were examined, as the PICA do not reach the cerebellum before P0. As illustrated in Fig. 8A, at E11, the SCA was confined to a discrete cerebellar domain, and the superficial vascular network did not yet fully cover the developing cerebellum. By E13, both the cerebellum and the SCA had undergone substantial development, characterized by the emergence of multiple arterial branches that increased vascular coverage across the cerebellar surface. At this stage, vessels were detected up to 300 µm from the SCA. However, this distal network exhibited large meshes. At E15, continued maturation of the SCA led to smaller and denser meshes in the lateral part of the cerebellar hemispheres (Fig. 8A). This resulted in a denser and more homogeneous vascular network throughout the cerebellum. A similar organization was maintained at E17. However, as the cerebellum expanded, the SCA became positioned more laterally, and vessels located within the prospective vermis appeared increasingly distant from the SCA branches. At P0, the marked increase in cerebellar volume expanded the surface area requiring vascularization. Despite extensive perpendicular branching of the SCA, several regions of the cerebellum remained located at relatively large distances from the arterial network (Fig. 8A). Regarding the AICA, observations at E15 and E17 revealed a progressive extension of the arteries toward the anteroinferior region of the cerebellum, although it remains at distance (Fig. 8B). By P0, the AICA established its first connections with the cerebellum and began to vascularize its inferior aspect (Fig. 8B).

**Fig. 8.**
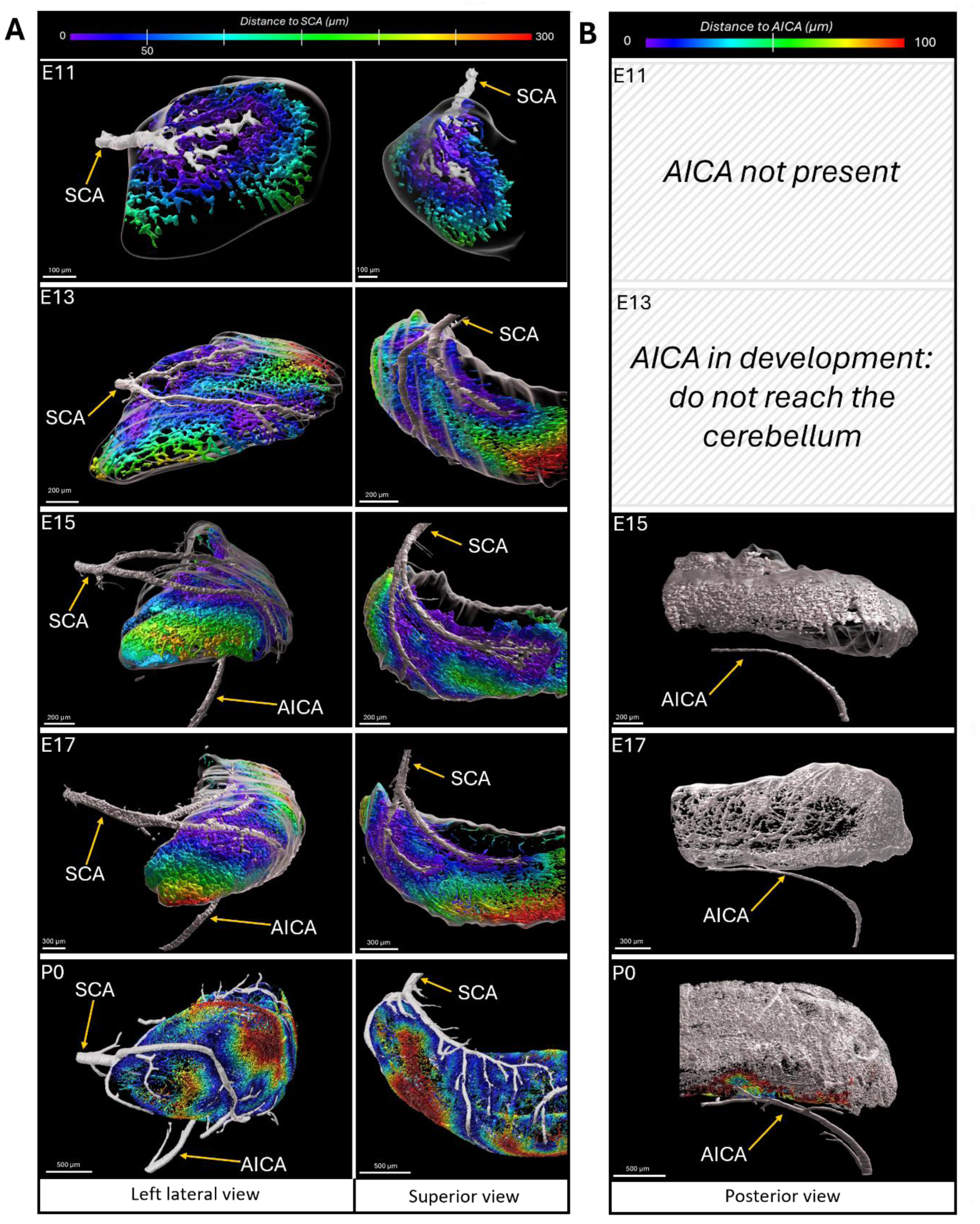
Architecture of the SCA and AICA contribution to the superficial vascular network of the cerebellum during the mouse embryonic development. (A) 3D modellings of the superficial vascular network of the cerebellum, viewed from above and in right side, illustrating the contribution of the SCA to the vascularization of the cerebellum in mouse embryos at E11, E13, E15, E17 and in a newborn at P0. Superficial vessels are color-coded according to their distance from the SCA within a range of 0–300 µm. (B) 3D modellings of the superficial vascular network of the cerebellum in a posterior view, illustrating the contribution of the AICA to the vascularization of the cerebellum in mouse embryos at E15, E17 and in a newborn at P0. Superficial vessels are color-coded according to their distance from the AICA within a range of 1–100 µm. AICA: anterior inferior cerebellar artery; SCA: superior cerebellar artery.

#### 3.4.2 Internal organization of the cerebellar vascularization

In order to model the vasculature of superficial and internal networks, the machine-learning assisted workflow we have developed on Imaris was applied (Rodriguez-Duboc et *al*., 2026). The use of this workflow to both networks enabled the reconstruction of the global vascular architecture of the cerebellum and its developmental evolution (Fig. 9).

**Fig. 9.**
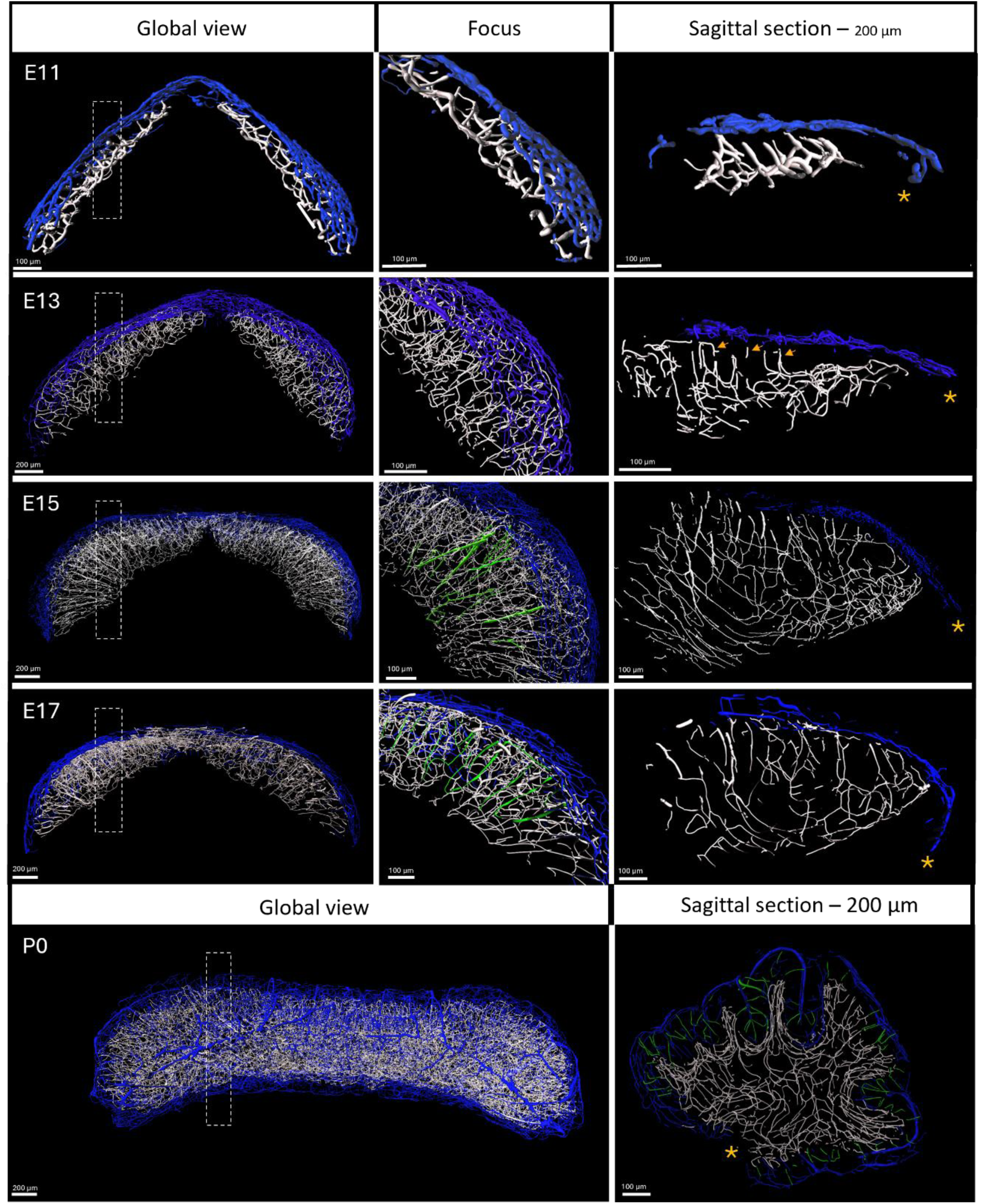
Evolution of the vascular architecture of the cerebellum in mouse over developmental stages. 3D modellings of the cerebellar vascular network in mouse embryos at E11, E13, E15, E17 and in a P0 newborn, illustrating the evolution of cerebellar vascularization during embryonic development. The superficial and internal vascular networks were modelled separately using filaments generated by machine-learning in Imaris and are shown in blue and white, respectively. An overview of the cerebellum is shown for each stage (on the right) with a higher magnified view of the right part for the embryonic stages (in the middle). To better visualize the internal organization of the vessels, a 200-µm thick sagittal section (represented by the dotted rectangle in the overview) is modelled on the right. The vessels colored in green represent the main radial vessels perpendicular to the superficial network. In E13, arrowheads indicate developing radial vessels. The choroid plexus is indicated by an asterisk in each sagittal section image.

At E11, the superficial and internal vascular networks were organized as two apposed vascular sheets. High magnification and sagittal sections reveal that vessels originating from the superficial network (blue network) penetrated radially into the cerebellar tissue, forming a scaffold that contributed to the establishment of the internal vascular layer (white network). At E13, the density of the intra-cerebellum vessels increased and displayed multidirectional orientation. However, radial penetrating vessels remained clearly identifiable and likely contributed to the progressive expansion of the vascular network within the core of the organ (Fig. 9, arrows). At E15, the internal vascular network underwent a further increase in volume and complexity, and extensively colonized the embryonic cerebellum. In parallel, the radial organization of penetrating vessels became prominent (Fig. 9, green vessels). At E17, the overall vascular organization remained similar to that observed at E15, although the radial vessels appeared thinner but more structured. By P0, during the onset of cerebellar lobulation, the superficial vascular network completely covered the cerebellar surface with regularly distributed radial vessels. These vessels gave rise to the entire internal vasculature, which displayed a more defined organization following the architecture of each developing lobule (Fig. 9, sagittal planes).

### 3.5 Morphometric parameters of the vasculature of the developing cerebellum

To quantitatively investigate the developmental evolution of the intra-cerebellar vascular network, we used a dedicated and highly detailed software, VesselVio, which is better suited for vascular parameter analysis.

Based on these parameters, a principal component analysis (PCA) was firstly performed (Fig. 10). The study of the variables show that the calculated dimensions represent for 73.9% of the total variance, with Dim1 explaining 47.4% and Dim2 explaining 26.5%, indicating that the Dim1–Dim2 factorial plane captures most of the complexity of the dataset. The correlation circle shows that Dim1 is mainly driven by variables reflecting global vascular network expansion and complexity (Fig. 10A). Vascular volume, surface area, number of segments, network length, branchpoints, endpoints, and cerebellar volume are all strongly associated with the positive side of this axis. In contrast, mean segment tortuosity, mean segment length, and segment partitioning contributes in a different direction and captures aspects of vascular organization that are less directly related to overall network size. Dim1 therefore represents a gradient of vascular maturation, ranging from small and less developed networks to larger, more extensive, and highly branched networks. Dim2 captures a second, more local dimension of vascular morphology. It is mainly associated with mean segment radius, mean segment volume, mean segment surface area, and vascular density, all of which contributed positively to this axis, whereas mean segment length contributes negatively. This indicates that Dim2 distinguishes networks composed of thicker vascular segments from those composed of longer and more elongated segments. Taken together, the two axes reflect complementary aspects of vascular development: one related to global expansion and branching, the other to segment-level morphology.

**Fig. 10.**
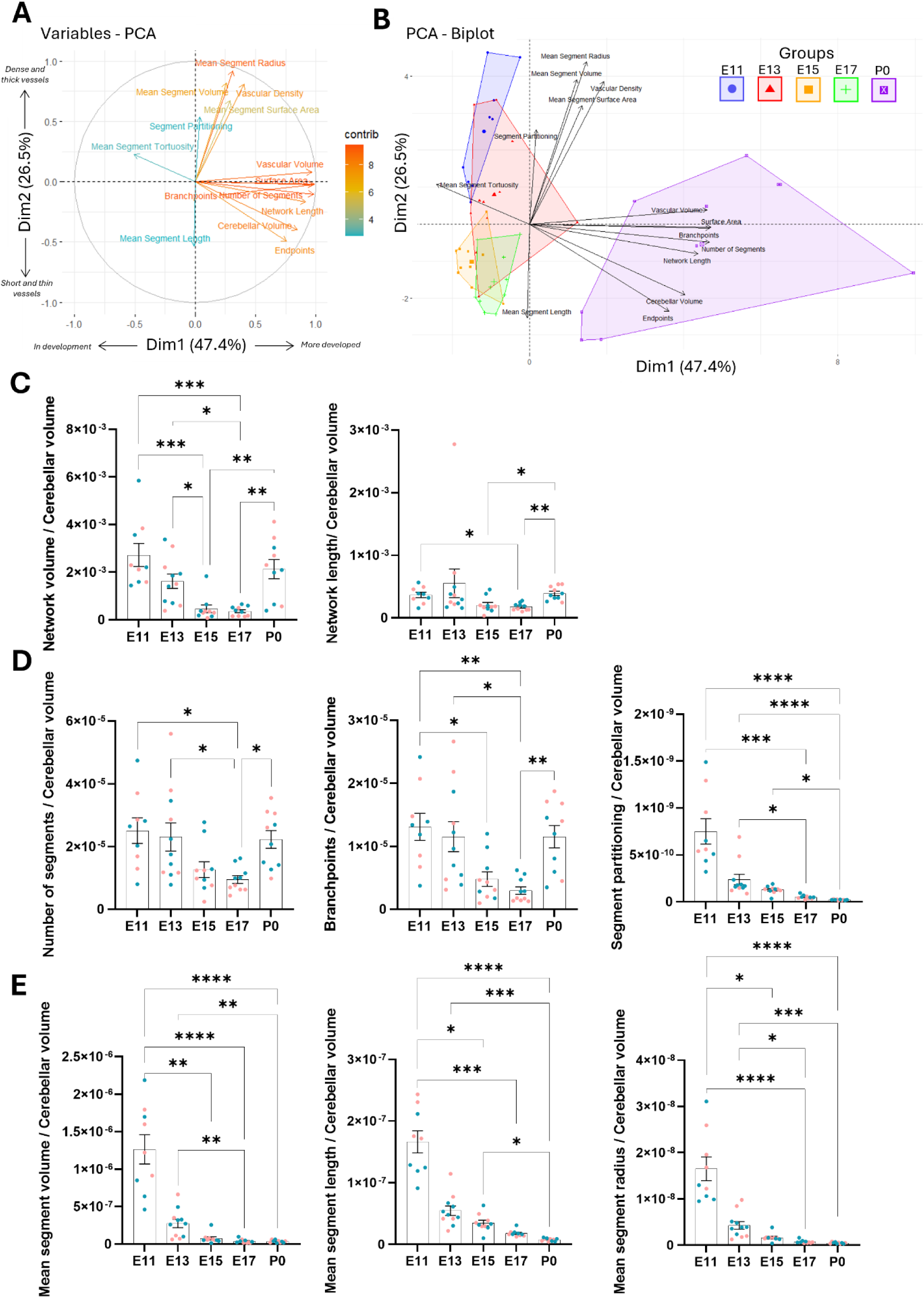
(next page) Analysis of the intra-cerebellar vasculature over mouse embryonic development. (A) Biplot representation of variables obtained from the quantitative analysis of vascular features. (B) Projection of individual samples onto the principal component analysis (PCA) factorial plane based on the 14 quantitative vascular parameters represented in A, illustrating the intra-cerebellar remodeling of the vascular network during mouse embryonic development. Each point represents an individual sample and is color-coded according to developmental stage (n = 10 per stage; 5 males and 5 females): E11 (yellow), E13 (green), E15 (blue), E17 (orange), and P0 (purple). (C-E) Quantification of global vascular architecture (C), overall segment composition (D) and mean segment morphology (E) in E11, E13, E15, E17 and P0 mice. Values are presented as mean ± SEM. Statistical analyses were performed using the Kruskal–Wallis test followed by Dunn’s post hoc test (*p-value < 0.05; **p-value < 0.01; ***p-value < 0.001; ****p-value < 0.0001). Individual data points are shown, with blue points representing males and red representing females.

Projection of the samples onto the factorial plane reveals a clear developmental progression from E11 to P0 (Fig. 10B). The embryonic groups are located mainly on the left side of the plot, close to the origin, and partially overlap with one another, indicating broadly similar and less differentiated vascular profiles. Among them, E11 is positioned in the upper-left region of the plane, in association with higher values of vascular density, mean segment radius, and mean segment volume. This indicates that this stage was characterized by a dense vascular network with thick segments, despite limited overall morphological complexity. E13 occupies an intermediate position partially overlapping both E11 and E15/E17, with more variable parameters, consistent with a transition between an immature dense network and a more expanded vascular organization. E15 and E17 are located closer to the lower-left quadrant and show a greater overlap, indicating that they belong to the same developmental trajectory. At these stages, the vascular network appeared to become more elongated, less dense, and less complex, consistent with ongoing remodeling and structural refinement. The overlap between E15 and E17 shows that the changes occurring between these two stages were gradual rather than abrupt. In contrast, the postnatal group P0 is clearly separated from the embryonic stages along the positive side of Dim1. This position reflects a marked increase in vascular volume, surface area, network length, number of segments, branchpoints, and endpoints, indicating a much more developed and complex vascular architecture. P0 therefore represented the most mature vascular state in the dataset, with an extensively branched network that differs markedly from the embryonic profiles.

To better characterize and clearly illustrate the evolution of cerebellar vascularization during embryonic development, histograms of the principal variables identified by the PCA were generated, distinguishing three major categories of parameters: global vascular architecture, overall segment composition, and mean segment morphology (Fig. 10C-E).

Regarding global vascular architecture, total vascular network volume showed a significant decrease from E11 to E17 (Kruskal–Wallis; 0.0006 < p-value < 0.0427), followed by a marked increase at P0 (Kruskal–Wallis; **p-value = 0.0056). Similarly, total network length was significantly reduced in E17 cerebella compared with E11 (Kruskal–Wallis; *p-value = 0.0244) and subsequently increased at P0 (Kruskal–Wallis; **p-value = 0.0033; Fig. 10C). In terms of overall vascular composition, the number of vascular segments remained relatively stable between E11 and E13, then progressively decreased until E17 (Kruskal–Wallis; *p-value = 0.0194), before increasing again at P0 (Kruskal–Wallis; **p-value = 0.0059). The number of branchpoints followed a similar developmental trajectory (Fig. 10D). Notably, segment partitioning exhibited a continuous decline throughout development, reaching its lowest value at P0 (Fig. 10D). Analysis of mean segment morphology further revealed significant reductions in mean segment volume, length, and radius up to birth (Fig. 10E). It should be noted that most vascular parameters are temporarily higher in males than in females at E17 (Table S4).

At P0, although total network length and segment number increased, mean segment volume remained reduced. In parallel, the decrease in segment partitioning suggests that vascular expansion outpaces network complexity, indicating a progressive structural organization and refinement of the vascular network (Fig. 10C).

## 4. Discussion

Despite significant progress in studying cerebellar neurogenesis, the understanding of its angiogenesis during embryogenesis remains limited.

Whole-embryo clearing, 3D imaging and automated segmentation enabled a detailed spatiotemporal characterization of cerebellar afferent vascular development from E11 to birth in the mouse. The 3D reconstruction reveals a sequential maturation of the hindbrain vasculature, initiated by the formation of a single basilar artery (BA) derived from the internal carotid arteries (ICA) at E11, followed by the emergence of the vertebral arteries (VA) at E13. This highly stereotyped sequence appears to be evolutionarily conserved across vertebrates, despite some exceptions, such as the presence of paired BA in dogfish for example (Rahmat and Gilland, 2013).

Then, cerebellar vascularization follows a coordinated developmental progression, characterized by the identification of the superior cerebellar arteries (SCA) coming from a primitive BA from E11 onwards. At E13, the transitional anastomoses have regressed, making it possible to clearly distinguish the BA, from which the AICA differentiate. These arteries reach the cerebellum at E17 and start to participate to the cerebellar vascularisation at P0. The situation regarding the PICA remains less well-defined. Emerging from the VA at E15 and progressing toward the cerebellum through P0, the PICA does not appear to reach the cerebellum in mouse during the embryonic period, and thus may not participate in cerebellar vascularization at least until birth. The contribution of the PICA to cerebellar blood supply remains controversial. Although it is widely accepted that cerebellar tissue receives arterial supply from three paired vessels—the SCA, AICA, and PICA—multiple studies conducted in primates indicate that only two arteries participate in cerebellar vascularization, thereby excluding the PICA from this contribution (Lake et *al*., 1990; Sabec-Pereira et *al.*, 2020). The fact that the emergence of PICA is associated with the development of the neocerebellum and therefore appears relatively late in mammalian phylogeny (Macchi et *al*., 2005) could explain the apparent inability to establish a clear topography of PICA in vertebrates. Nevertheless, some anatomical landmarks make possible to compare the time course of the cerebellar arteries in mice and humans during development. Thus, at E11, the emergence of the primitive SCA and their progressive encirclement of the cerebellum, makes this mouse stage comparable to the Padget stages 3 (7–12 mm; ≃ 33 days) and 4 (12–14 mm stage; ≃ 36 days) in human embryos. Then, the progressive emergence of the AICA and the PICA as primitive vessels at E13 closely corresponds to human Padget stage 5 (16–18 mm; ≃ 40 days). At E15, the continued development of the AICA and PICA, without reaching the cerebellar surface, is comparable to the vascular configuration observed at Padget stage 6 (20–24 mm; ≃ 45 days). At E17, the anterior convergence of the AICA toward the cerebellum parallels the vascular arrangement described at Padget stage 7 (40 mm stage; ≃ 50 days). Finally, at P0, the PICA are well developed, exhibiting numerous collateral branches, and may mark the transition between the embryonic and fetal patterns of cerebellar vascularization (Bertulli and Robert, 2021).

The 3D reconstruction also permits quantitative analysis of morphometric evolution of these arteries. This revealed a quasi-constant SCA-to-cerebellar volume ratio throughout development, whereas AICA and PICA volumes increased over time. These findings suggest that the SCA follow a developmental trajectory tightly coupled to cerebellar morphogenesis. This is consistent with their early emergence and their conserved role as the primary arteries dedicated to cerebellar vascularization (Macchi et *al*., 2005). In contrast, the later emergence and faster growth of the AICA and PICA until E15 may reflect their broader territorial functions, as both arteries contribute to additional hindbrain structures beyond the cerebellum (Savoiardo et *al*., 1987). Thus, their more rapid morphometric progression may be linked to the progressive acquisition of multiple vascular territories rather than to cerebellar growth alone (Delion et *al*., 2016). The decrease in the artery length-to-cerebellar volume ratio suggests that at E17 these arteries do not need to elongate further to match cerebellar growth.

By reassigning the data to individual embryos, it becomes possible to reveal both intra- and inter-individual heterogeneity. As expected, significant variability is observed among the different cerebellar artery types. More importantly, this analysis shows that, within a given arterial pair, the developmental trajectory could differ between the left and right sides, as illustrated by the AICA at E15. Furthermore, marked inter-individual variability emerges from E13 onward, together with a sex-related effect on the SCA at E11 and on the right branches of all cerebellar arteries at E15.

Such heterogeneity is consistent with the diversity of cerebellar arterial topography that has been well described in humans. Indeed, our results show that, as in humans, the mouse SCA display a relatively homogeneous organization, whereas the AICA and PICA exhibit considerable variability in their origin (Ballesteros-Acuña et *al*., 2022; Scotti, 1975), morphology (Macchi et *al.*, 2005; Sharifi and Ciszek, 2010), and branching patterns (Goto and Inoue, 2022).Overall, these findings suggest that mouse cerebellar arteries retain a degree of morphological plasticity comparable to that reported in human posterior circulation (Błaszczyk et *al.*, 2024).

Together, these arteries initiate vascularization of the cerebellar surface before extending into the cerebellar parenchyma, although their respective contributions differ markedly. The SCA are the first and principal arteries to participate in this process. They initially establish a superficial vascular network over the cerebellar surface through a medial branch that courses around the superior aspect of the organ and a lateral branch that extends along its lateral surface toward the midline, corresponding to the future vermis. At the earliest stages, the vascular meshwork is characterized by large meshes and low vessel density. As development proceeds, the meshes progressively narrow while the vascular density increases, resulting in an almost complete coverage of the cerebellar surface by P0. This developmental sequence is consistent with the classical mechanisms of angiogenesis. It begins with an initial sprouting phase, which generates a sparse vascular network with relatively large meshes, followed by a phase of intussusceptive angiogenesis that rapidly expands and remodels the network through intraluminal vessel splitting (Djonov et *al*., 2003; Nitzsche et *al*., 2022). In contrast, the AICA contribute at a later stage, progressively extending toward the anteroinferior surface of the cerebellum and initiating vascularization of this territory around birth. The contribution of the PICA to cerebellar vascularization is less clear. Although designated as the posterior inferior cerebellar arteries, our data indicate that, in mice, the PICA do not reach the cerebellum before birth. Their contribution may therefore be exclusively postnatal, or they may participate indirectly in early cerebellar vascularization by branching to other arteries such as BA or AICA for example. This organization may be interpreted in the context of the AICA–PICA dominance concept, which proposes that the size of the AICA and the extent of its vascular territory are inversely related to those of the PICA, and vice versa (Lasjaunias et *al*., 2011). It has been also reported that the PICA give rise to choroidal branches contributing to the vascularization of the fourth ventricular choroid plexus (Fujii et *al*., 1980; Sharifi et *al*., 2005). This suggests that its early developmental role may be primarily associated with the choroidal and periventricular vascular territories rather than the cerebellar parenchyma. Taken together, these observations indicate that, during mouse embryonic development, the AICA predominantly supply both the anterior and posterior inferior regions of the cerebellum, and the PICA may preferentially contribute to the vascularization of the myelencephalic plexus before eventually establishing its definitive cerebellar territory postnatally.

The internal cerebellar vasculature progressively arises from the superficial vascular network, most likely as part of the general program of CNS angiogenesis and following the developmental sequence described by Fantin et al. (2013). Radial vessels first emerge from the superficial arterial meshwork, which can be considered analogous to the perineural vascular plexus (PNVP), and penetrate the cerebellar parenchyma toward the ventricular zone. They subsequently bend and anastomose to establish a ventricular vascular plexus by E11. From this stage onward, lateral sprouts emerge and progressively interconnect, giving rise to an increasing vascular network. However, the expansion of the intracerebellar vascular network does not initially keep pace cerebellar growth, resulting in a decline in the ratio vascular length/cerebellar volume until E17. This trend is accompanied by a reduction in segment number, segment length, segment diameter, and branchpoint number, suggesting a transient simplification of the vascular architecture as the network adapts to the rapid expansion of the cerebellum. Between E15 and E17, although some segment parameters exhibit transient sex-dependent differences, most of them remain stable, indicating that the vascular growth proceeds proportionally with cerebellar expansion. Nevertheless, at E17, vascular modelling becomes apparent with thinning of the radial vessels and reduced interconnectivity, leading to the emergence of less vascularized territories that foreshadow the future cerebellar lobules. At P0, vascularization undergoes a marked acceleration, resulting in increased vessel density as well as higher numbers of vascular segments and branchpoints. Radial vessels extend through the molecular layer and give rise to a well-developed vascular network throughout the cerebellar parenchyma. Consequently, the vascular architecture becomes both more extensive and more hierarchically organized, following the developing lobular pattern and explaining the decrease in network partitioning. This organization is maintained and continues to mature throughout postnatal development (De Launoit et *al*., 2026; Rodriguez-Duboc et *al*., 2026). Interestingly, the superficial vascular network does not extend into the cerebellar parenchyma at the lateral region corresponding to the former rhombic lip. One possible explanation is the close proximity of the myelencephalic choroid plexus, which originates from this region and becomes highly vascularized at an early embryonic stage. The presence of this pre-existing vascular bed may reduce the requirement for direct invasion by the superficial cerebellar vascular network (Macchi et *al*., 2005).

With respect to neurogenesis, the successive stages of vascularization can be related to the emergence of the different cerebellar neuronal populations and structures derived from the two embryonic germinal zones. At E11, when the first superficial vascular sheet is established, the ventricular zone is already well defined and constitutes a highly proliferative domain with substantial metabolic demands (Haldipur et *al*., 2019). This germinal zone sequentially gives rise to radial glial cells, GABAergic neurons of the cerebellar nuclei, Purkinje cells, and, slightly later, Golgi cells and cortical GABAergic interneurons (Leto et *al*., 2016). Neuronal differentiation then progressively extends above the ventricular zone between E11 and E15, concomitantly with the expansion of the intra-cerebellar vascular network. During this period, blood vessels are likely to support cerebellar maturation by providing oxygen and metabolic substrates. Indeed, during postnatal development, hypoxia has been shown to maintain granule cell precursors in a proliferative state (Kullmann et *al*., 2020), suggesting that the progressive increase in vascularization during embryogenesis may promote neuronal differentiation by improving tissue oxygenation. As the vascular network becomes more organized from E17 onward, it may also provide a structural scaffold for neuronal migration, as previously demonstrated for GABAergic neurons in the retina and cerebral cortex (Dumanoir et *al*., 2023; Won et *al*., 2013).

The rhombic lip constitutes the second germinal zone of the cerebellum. It remains highly proliferative between E11 and E17 and generates glutamatergic neurons of the cerebellar nuclei between E10.5 and E12.5, followed by granule cell precursors from E13.5 onward (Haldipur et *al*., 2019; Leto et *al*., 2016). During this embryonic period, the rhombic lip is primarily vascularized by the myelencephalic vascular plexus. However, our results indicate that the cerebellar arteries also contribute to its vascular supply along its external surface. This dual vascularization may contribute to the molecular regionalization described in the murine rhombic lip (Haldipur et *al*., 2019). One hypothesis is that granule cell precursors located adjacent to the superficial vascular network preferentially migrate along this vascular bed to establish the external granule cell layer, whereas progenitors located deeper and in closer proximity to the myelencephalic vascular plexus may preferentially contribute to the posterior vermis.

In this study, we provide the first comprehensive characterization of cerebellar vascular development during mouse embryogenesis. By establishing a developmental atlas of cerebellar vascularization, our work provides a foundation for future studies investigating the role of vascular development in cerebellar maturation and its contribution to perinatal vascular injury, such as hemorrhage or infarction (Pierson and Al Sufiani, 2016), and neurodevelopmental disorders, including autism (Azmitia et *al*., 2016).

## Declarations

### Data and Code Availability Statement

The R code used in this study is described in the Materials and Methods section. Statistical analyses and corresponding p-values are reported in supplementary data.

### Declaration of generative AI and AI-assisted technologies in the manuscript preparation process

ChatGPT was used to correct grammatical errors and improve the flow of the English text, and the machine learning service of the Imaris software was used for training in vascular modelling. The authors reviewed and edited the output as needed and take full responsibility for the content of the published article.

### CRediT author statement

**Camille Racine:** Investigation, Methodology, Project administration, Data Curation, Formal analysis, Visualization, Validation, Writing - Original Draft, Writing - Review and Editing. **Bruno Gonzalez:** Funding acquisition, Resources, Writing - Review and Editing. **Delphine Burel:** Investigation, Conceptualization, Methodology, Funding acquisition, Project administration, Supervision, Resources, Writing - Original Draft, Writing - Review and Editing.

## Acknowledgments

We acknowledge the France-BioImaging infrastructure (https://ror.org/01y7vt929) supported by the French National Research Agency (ANR-24-INBS-0005 FBI BIOGEN). We are also grateful to Clément Grosol, Lucien Perran and Marvin Bidoda Seke, Master’s students in ‘Cellular Imaging’ at the University of Rouen Normandie, for their help with the acquisition and analysis of the 3D images. We would like to thank Christophe Chamot for his help with the 3D image acquisition and Alexis Lebon for his help with the software. Finally, we are really grateful to Garris Rebin, animal caretaker in the Biological Resources Department of the Faculty of Sciences at the University of Rouen Normandy, for his invaluable assistance with mouse breeding.

## Supplementary data

**Supplementary table S1:**
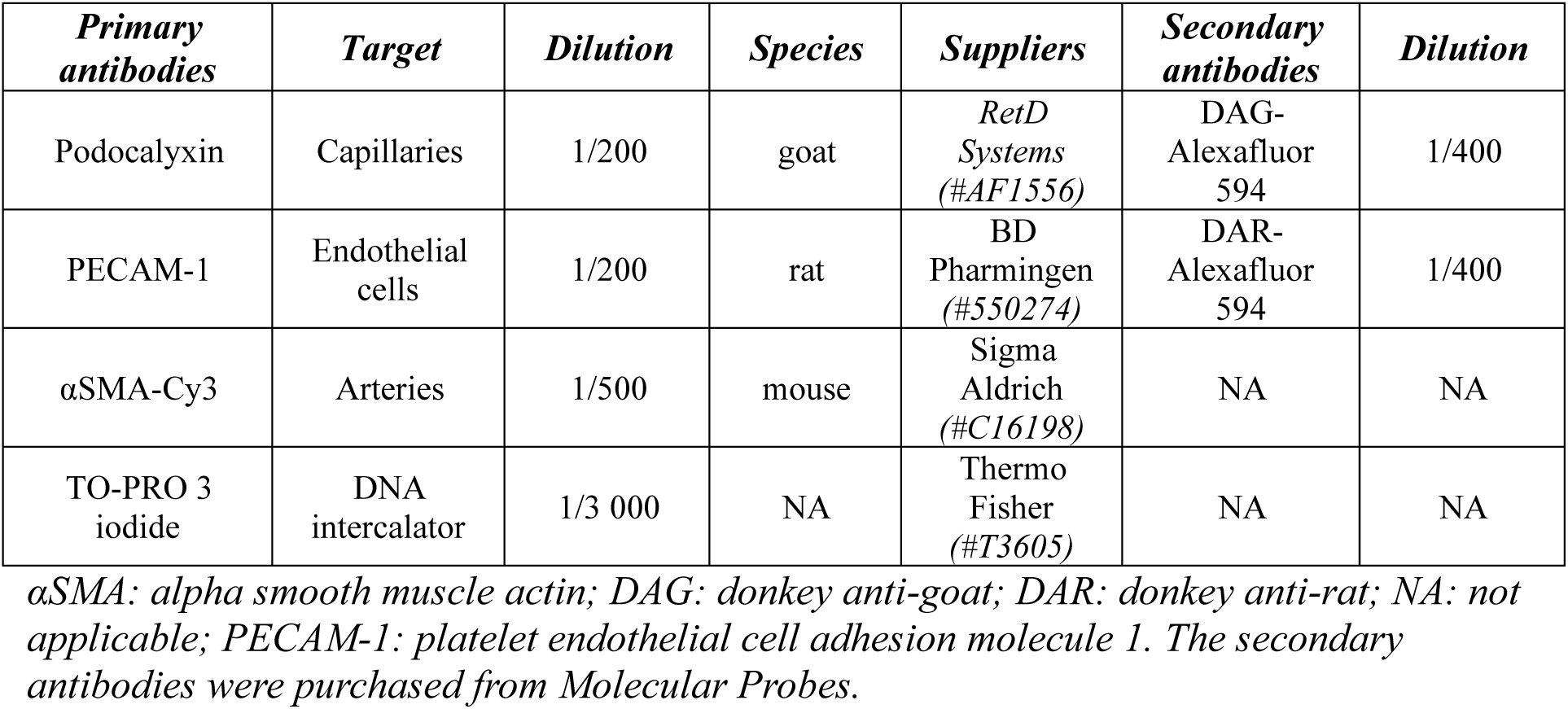
Antibodies and markers used for the visualization of blood vessels and cells respectively.

**Supplementary table S2:**
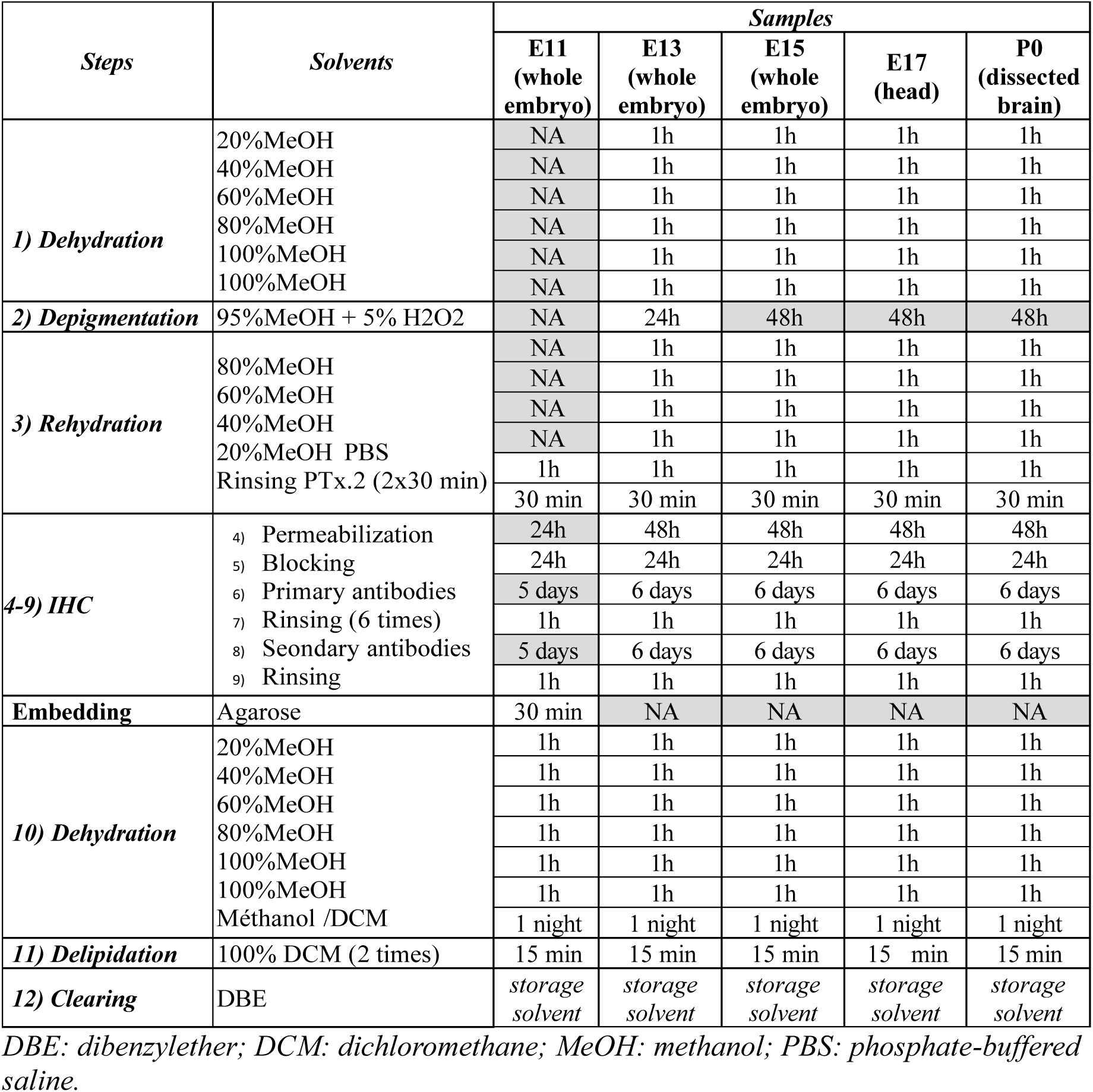
Adaptation of the iDISCO protocol depending on the age of the embryos/pups.

| *Steps* | *Solvents* | *Samples* |  |  |  |  |
| --- | --- | --- | --- | --- | --- | --- |
|  |  | **E11**<br>**(whole embryo)** | **E13**<br>**(whole embryo)** | **E15**<br>**(whole embryo)** | **E17**<br>**(head)** | **P0**<br>**(dissected brain)** |
| **1) Dehydration** | 20%MeOH | NA | 1h | 1h | 1h | 1h |
|  | 40%MeOH | NA | 1h | 1h | 1h | 1h |
|  | 60%MeOH | NA | 1h | 1h | 1h | 1h |
|  | 80%MeOH | NA | 1h | 1h | 1h | 1h |
|  | 100%MeOH | NA | 1h | 1h | 1h | 1h |
|  | 100%MeOH | NA | 1h | 1h | 1h | 1h |
| **2) Depigmentation** | 95%MeOH + 5% H2O2 | NA | 24h | 48h | 48h | 48h |
| **3) Rehydration** | 80%MeOH | NA | 1h | 1h | 1h | 1h |
|  | 60%MeOH | NA | 1h | 1h | 1h | 1h |
|  | 40%MeOH | NA | 1h | 1h | 1h | 1h |
|  | 20%MeOH PBS | NA | 1h | 1h | 1h | 1h |
|  | Rinsing PTx.2 (2x30 min) | 1h | 1h | 1h | 1h | 1h |
|  |  | 30 min | 30 min | 30 min | 30 min | 30 min |
| **4-9) IHC** | 4) Permeabilization | 24h | 48h | 48h | 48h | 48h |
|  | 5) Blocking | 24h | 24h | 24h | 24h | 24h |
|  | 6) Primary antibodies | 5 days | 6 days | 6 days | 6 days | 6 days |
|  | 7) Rinsing (6 times) | 1h | 1h | 1h | 1h | 1h |
|  | 8) Secondary antibodies | 5 days | 6 days | 6 days | 6 days | 6 days |
|  | 9) Rinsing | 1h | 1h | 1h | 1h | 1h |
| **Embedding** | Agarose | 30 min | NA | NA | NA | NA |
| **10) Dehydration** | 20%MeOH | 1h | 1h | 1h | 1h | 1h |
|  | 40%MeOH | 1h | 1h | 1h | 1h | 1h |
|  | 60%MeOH | 1h | 1h | 1h | 1h | 1h |
|  | 80%MeOH | 1h | 1h | 1h | 1h | 1h |
|  | 100%MeOH | 1h | 1h | 1h | 1h | 1h |
|  | 100%MeOH | 1h | 1h | 1h | 1h | 1h |
|  | Méthanol /DCM | 1 night | 1 night | 1 night | 1 night | 1 night |
| **11) Delipidation** | 100% DCM (2 times) | 15 min | 15 min | 15 min | 15 min | 15 min |
| **12) Clearing** | DBE | storage solvent | storage solvent | storage solvent | storage solvent | storage solvent |

DBE: dibenzylether; DCM: dichloromethane; MeOH: methanol; PBS: phosphate-buffered saline.

**Supplementary table S3:**
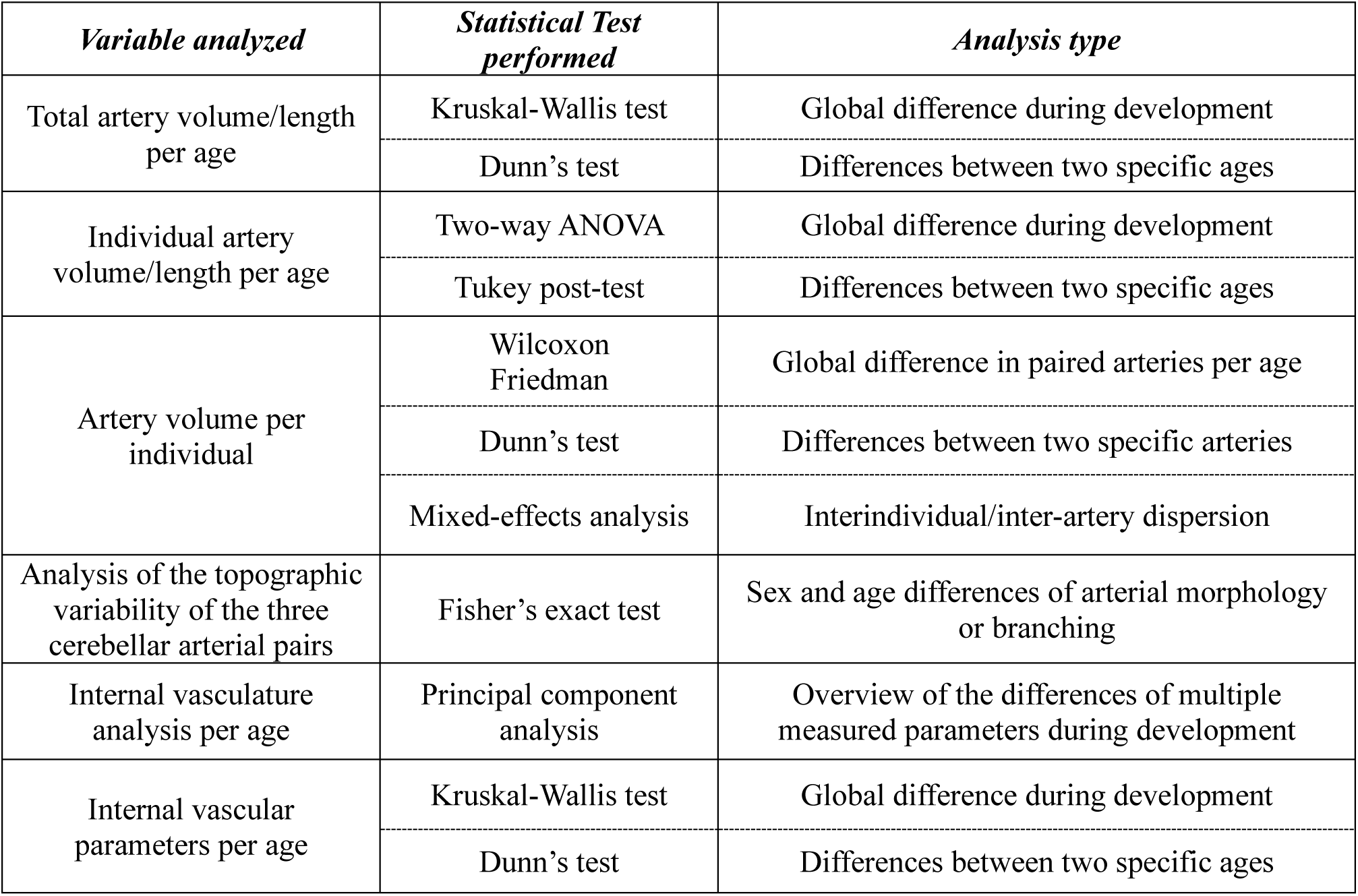
Summary of the statistical methods used for the analysis of the different variable types.

**Supplementary table S4:**
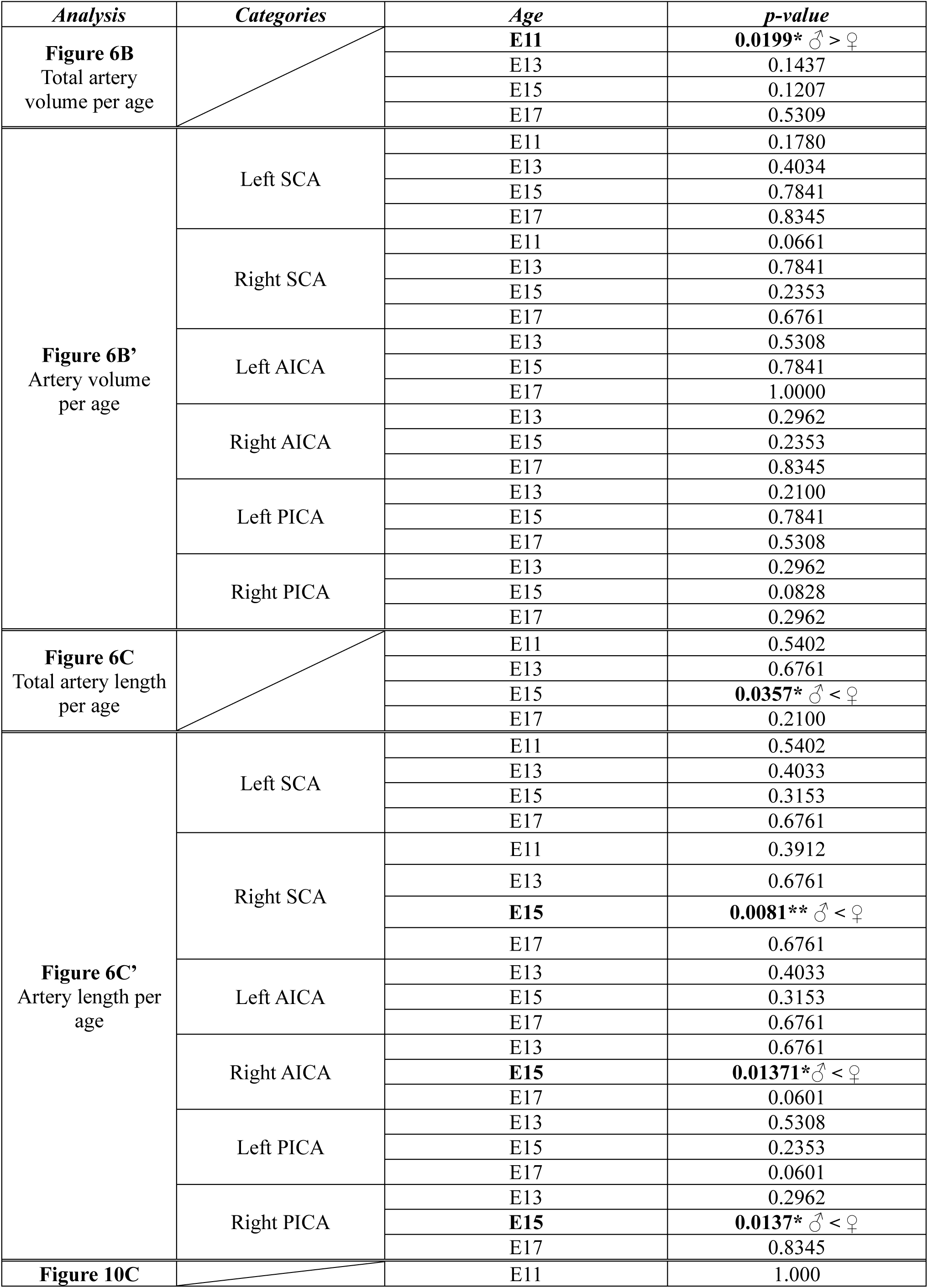
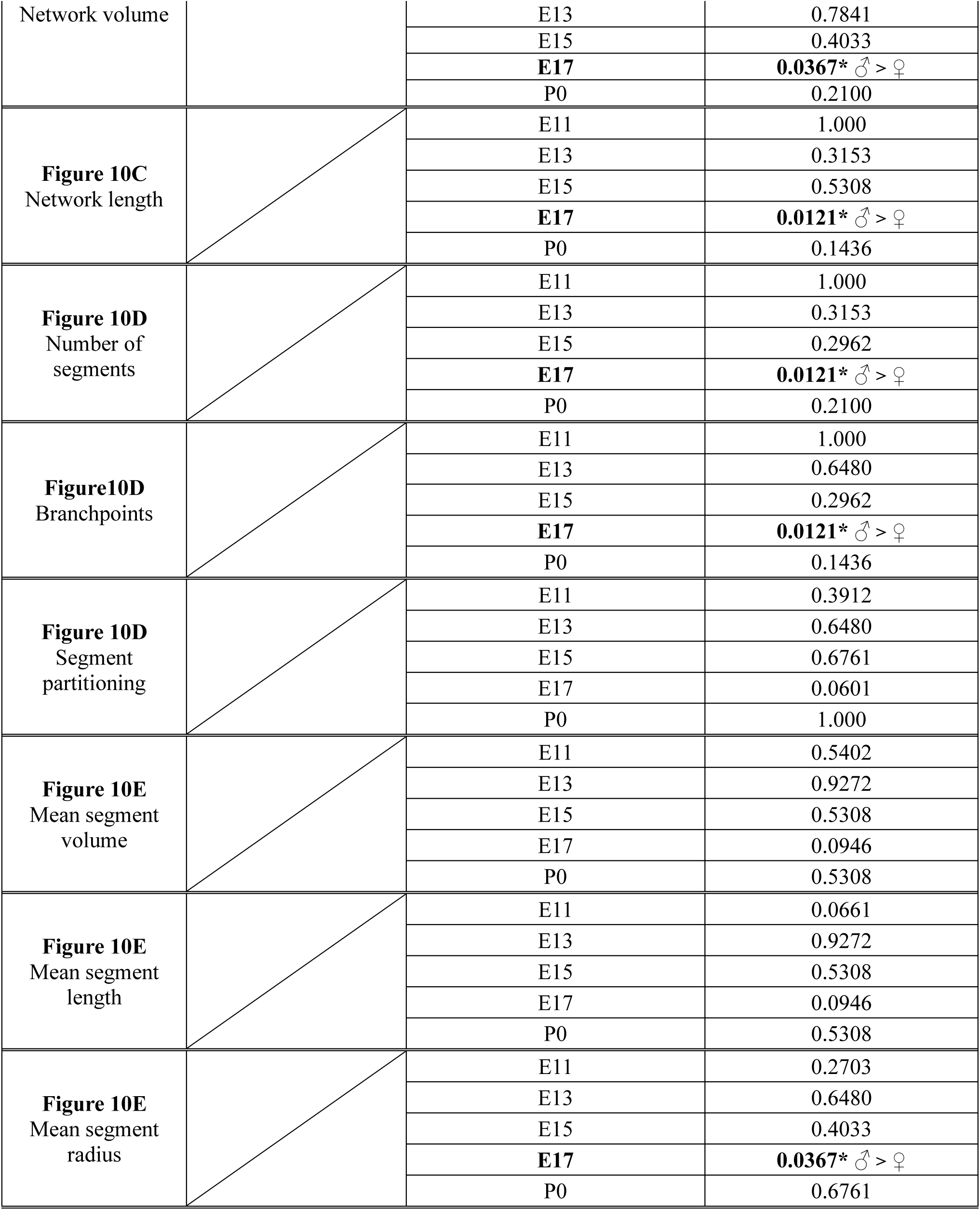
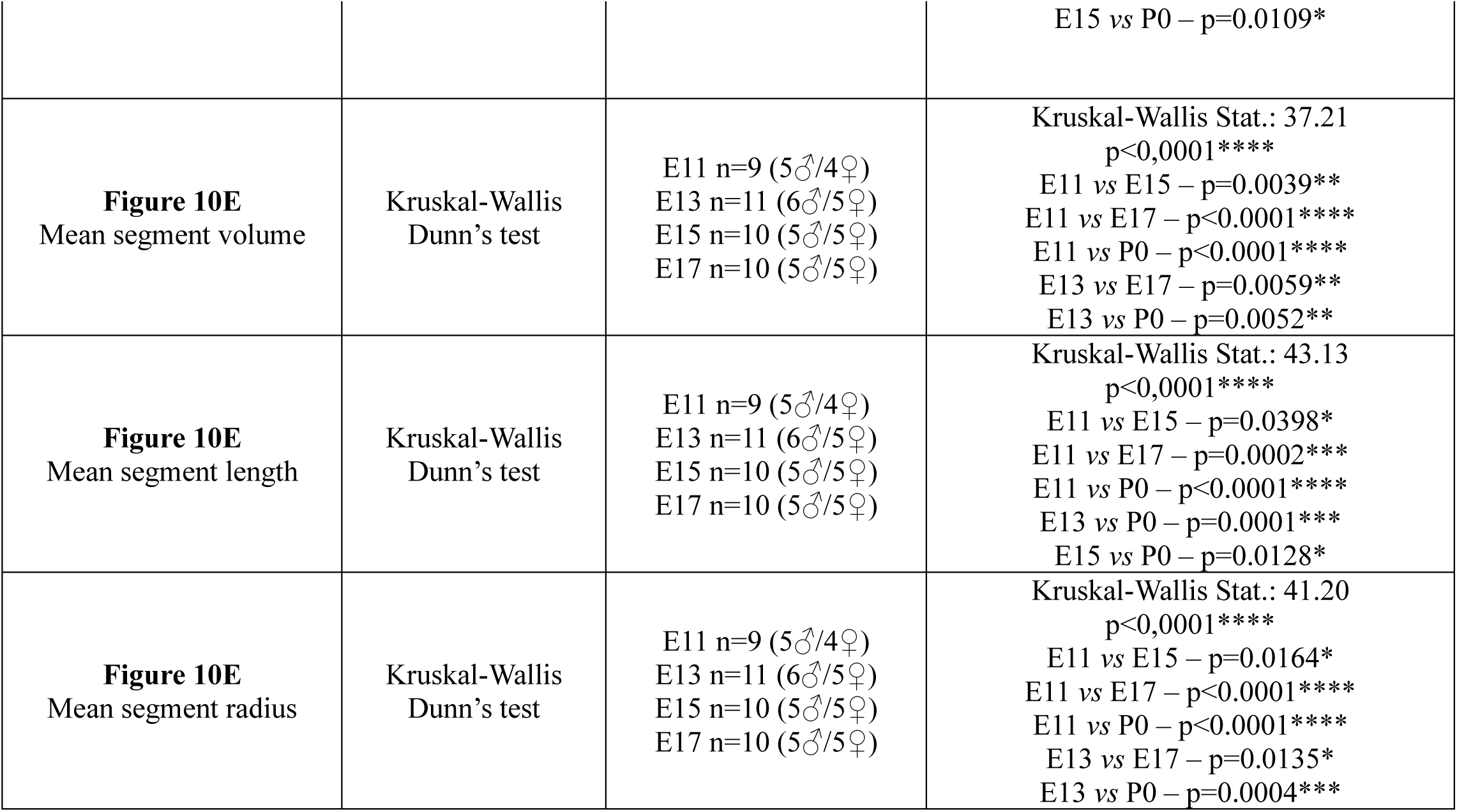
Sex effect analyses assessed using the Wilcoxon test. Variables with a significant sex effect (Wilcoxon rank-sum test, p-value < 0.05) are shown in bold.

**Supplementary table S5:**
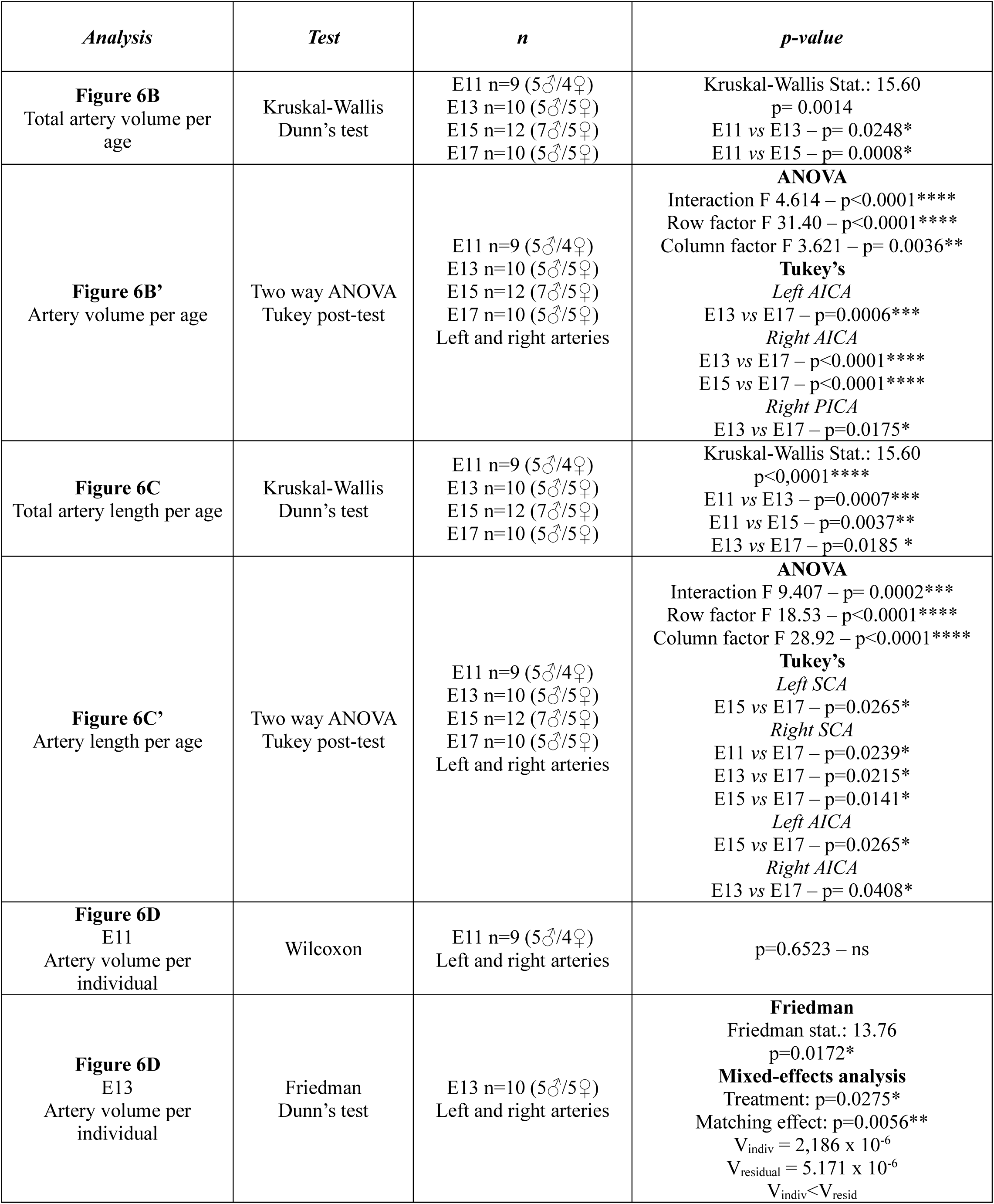
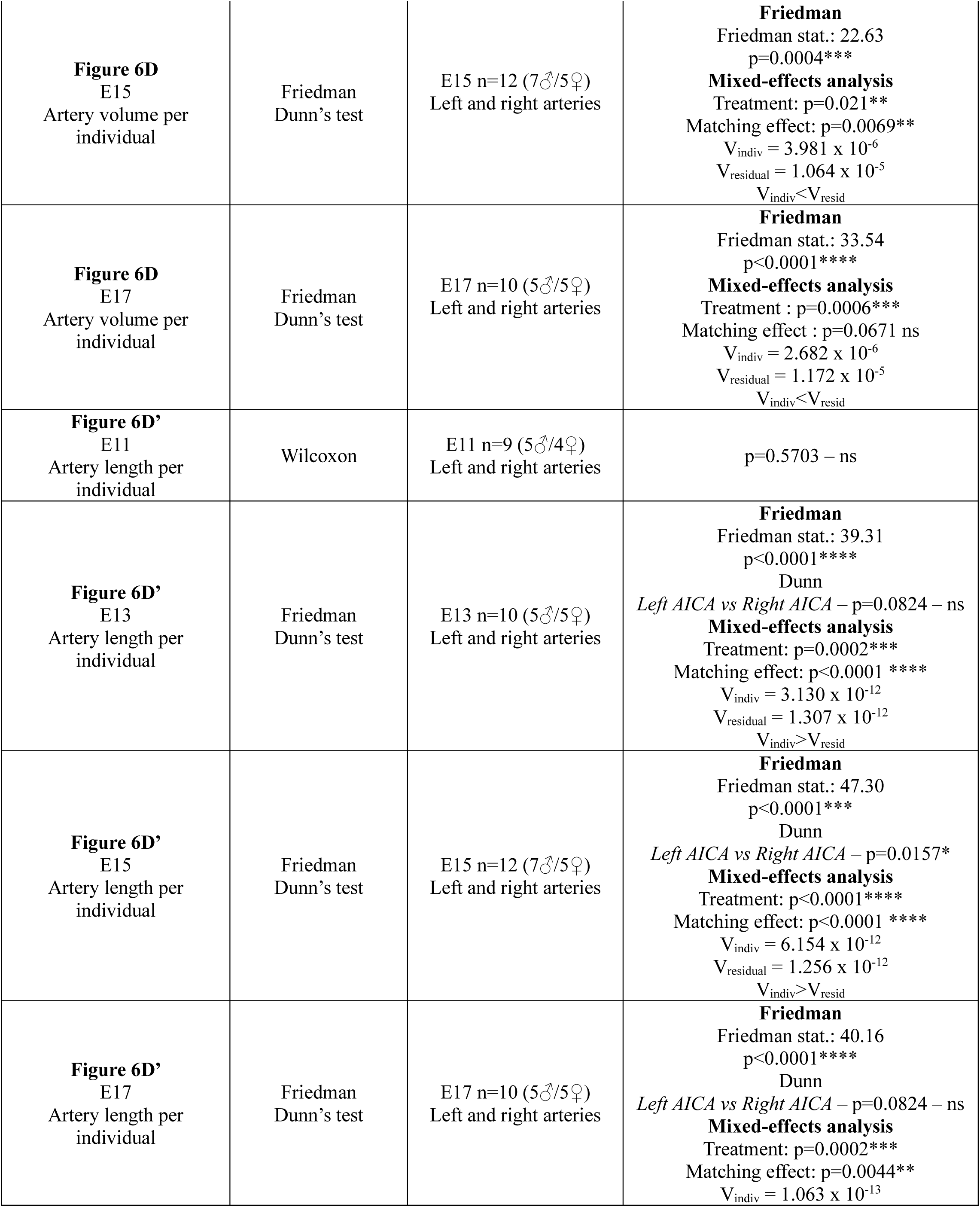
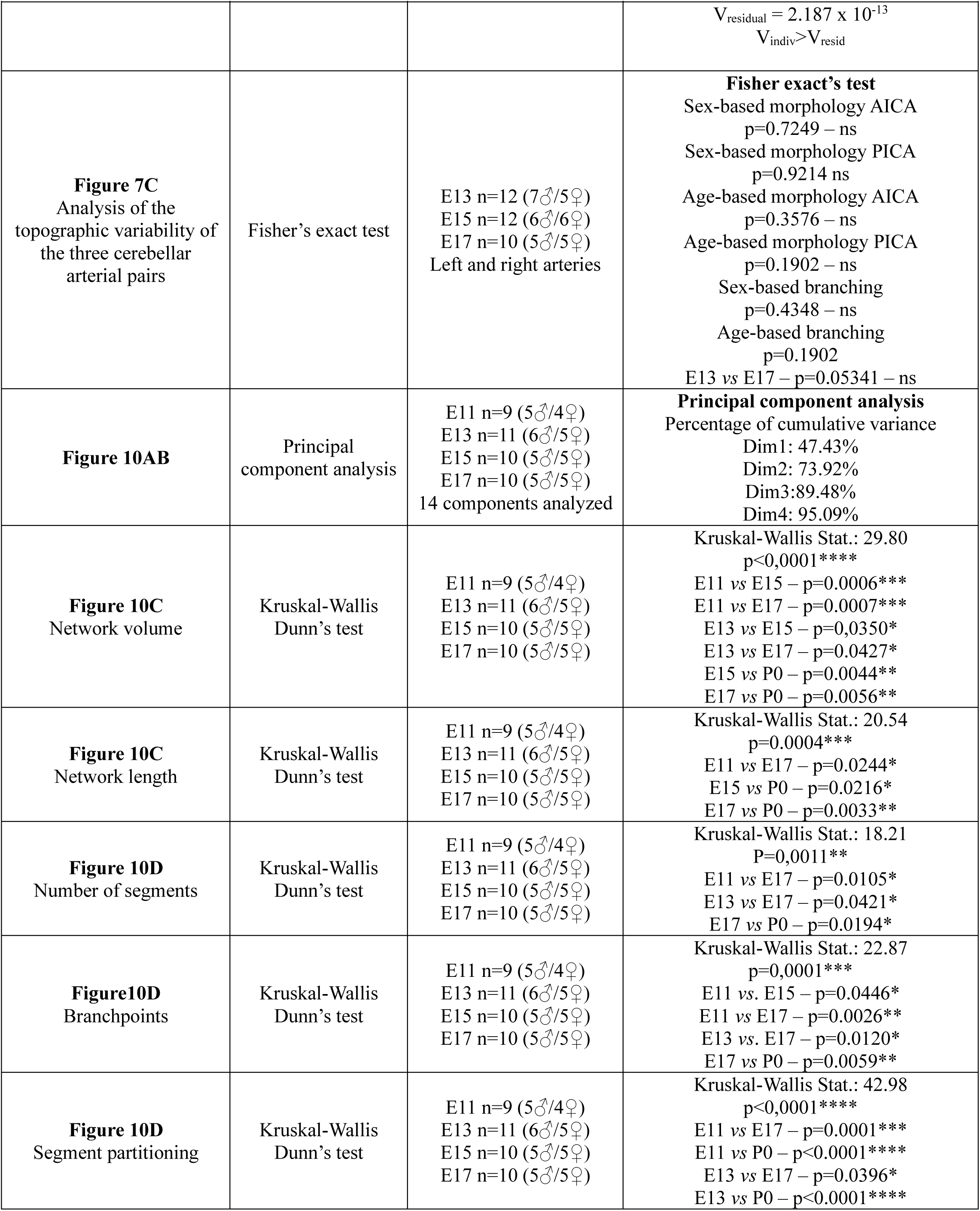
Summary of the statistical analyses used for each quantitative parameter.

