## Supplementary material for "Spatio-temporal 3D Mapping of Mouse Cerebellar Vascularization during Embryonic Development": Table S

**Supplementary data**

**Supplementary table S1: Antibodies and markers used for the visualization of blood vessels and cells respectively.**

| ***Primary antibodies*** | ***Target*** | ***Dilution*** | ***Species*** | ***Suppliers*** | ***Secondary antibodies*** | ***Dilution*** |
| --- | --- | --- | --- | --- | --- | --- |
| Podocalyxin | Capillaries | 1/200 | goat | *RetD Systems (#AF1556)* | DAG-Alexafluor 594 | 1/400 |
| PECAM-1 | Endothelial cells | 1/200 | rat | BD Pharmingen *(#550274)* | DAR-Alexafluor 594 | 1/400 |
| αSMA-Cy3 | Arteries | 1/500 | mouse | Sigma Aldrich *(#C16198)* | NA | NA |
| TO-PRO 3 iodide | DNA intercalator | 1/3 000 | NA | Thermo Fisher *(#T3605)* | NA | NA |

*αSMA: alpha smooth muscle actin; DAG: donkey anti-goat; DAR: donkey anti-rat; NA: not applicable; PECAM-1: platelet endothelial cell adhesion molecule 1.* *The secondary antibodies were purchased from Molecular Probes.*

**Supplementary table S2: Adaptation of the iDISCO protocol depending on the age of the embryos/pups.**

| ***Steps*** | ***Solvents*** | ***Samples*** | | | | |
| --- | --- | --- | --- | --- | --- | --- |
|  |  | **E11**  **(whole embryo)** | **E13**  **(whole embryo)** | **E15**  **(whole embryo)** | **E17**  **(head)** | **P0**  **(dissected brain)** |
| ***1) Dehydration*** | 20%MeOH  40%MeOH  60%MeOH  80%MeOH  100%MeOH  100%MeOH | NA | 1h | 1h | 1h | 1h |
|  |  | NA | 1h | 1h | 1h | 1h |
|  |  | NA | 1h | 1h | 1h | 1h |
|  |  | NA | 1h | 1h | 1h | 1h |
|  |  | NA | 1h | 1h | 1h | 1h |
|  |  | NA | 1h | 1h | 1h | 1h |
| ***2) Depigmentation*** | 95%MeOH + 5% H2O2 | NA | 24h | 48h | 48h | 48h |
| ***3) Rehydration*** | 80%MeOH  60%MeOH  40%MeOH  20%MeOH PBS  Rinsing PTx.2 (2x30 min) | NA | 1h | 1h | 1h | 1h |
|  |  | NA | 1h | 1h | 1h | 1h |
|  |  | NA | 1h | 1h | 1h | 1h |
|  |  | NA | 1h | 1h | 1h | 1h |
|  |  | 1h | 1h | 1h | 1h | 1h |
|  |  | 30 min | 30 min | 30 min | 30 min | 30 min |
| ***4-9) IHC*** | 4) Permeabilization 5) Blocking 6) Primary antibodies 7) Rinsing (6 times) 8) Seondary antibodies 9) Rinsing | 24h | 48h | 48h | 48h | 48h |
|  |  | 24h | 24h | 24h | 24h | 24h |
|  |  | 5 days | 6 days | 6 days | 6 days | 6 days |
|  |  | 1h | 1h | 1h | 1h | 1h |
|  |  | 5 days | 6 days | 6 days | 6 days | 6 days |
|  |  | 1h | 1h | 1h | 1h | 1h |
| **Embedding** | Agarose | 30 min | NA | NA | NA | NA |
| ***10) Dehydration*** | 20%MeOH  40%MeOH  60%MeOH  80%MeOH  100%MeOH  100%MeOH  Méthanol /DCM | 1h | 1h | 1h | 1h | 1h |
|  |  | 1h | 1h | 1h | 1h | 1h |
|  |  | 1h | 1h | 1h | 1h | 1h |
|  |  | 1h | 1h | 1h | 1h | 1h |
|  |  | 1h | 1h | 1h | 1h | 1h |
|  |  | 1h | 1h | 1h | 1h | 1h |
|  |  | 1 night | 1 night | 1 night | 1 night | 1 night |
| ***11) Delipidation*** | 100% DCM (2 times) | 15 min | 15 min | 15 min | 15 min | 15 min |
| ***12) Clearing*** | DBE | *storage solvent* | *storage solvent* | *storage solvent* | *storage solvent* | *storage solvent* |

*DBE: dibenzylether; DCM: dichloromethane; MeOH: methanol; PBS: phosphate-buffered saline.*

**Supplementary table S3:** **Summary of the statistical methods used for the analysis of the different variable types.**

| ***Variable analyzed*** | ***Statistical Test performed*** | ***Analysis type*** |
| --- | --- | --- |
| Total artery volume/length per age | Kruskal-Wallis test | Global difference during development |
|  | Dunn’s test | Differences between two specific ages |
| Individual artery volume/length per age | Two-way ANOVA | Global difference during development |
|  | Tukey post-test | Differences between two specific ages |
| Artery volume per individual | Wilcoxon  Friedman | Global difference in paired arteries per age |
|  | Dunn’s test | Differences between two specific arteries |
|  | Mixed-effects analysis | Interindividual/inter-artery dispersion |
| Analysis of the topographic variability of the three cerebellar arterial pairs | Fisher’s exact test | Sex and age differences of arterial morphology or branching |
| Internal vasculature analysis per age | Principal component analysis | Overview of the differences of multiple measured parameters during development |
| Internal vascular parameters per age | Kruskal-Wallis test | Global difference during development |
|  | Dunn’s test | Differences between two specific ages |

**Supplementary table S4: Sex effect analyses assessed using the Wilcoxon test.** Variables with a significant sex effect (Wilcoxon rank-sum test, p-value < 0.05) are shown in bold.

| ***Analysis*** | ***Categories*** | ***Age*** | ***p-value*** |
| --- | --- | --- | --- |
| **Figure 6B**  Total artery volume per age |  | **E11** | **0.0199* ♂ > ♀** |
|  |  | E13 | 0.1437 |
|  |  | E15 | 0.1207 |
|  |  | E17 | 0.5309 |
| **Figure 6B’**  Artery volume per age | Left SCA | E11 | 0.1780 |
|  |  | E13 | 0.4034 |
|  |  | E15 | 0.7841 |
|  |  | E17 | 0.8345 |
|  | Right SCA | E11 | 0.0661 |
|  |  | E13 | 0.7841 |
|  |  | E15 | 0.2353 |
|  |  | E17 | 0.6761 |
|  | Left AICA | E13 | 0.5308 |
|  |  | E15 | 0.7841 |
|  |  | E17 | 1.0000 |
|  | Right AICA | E13 | 0.2962 |
|  |  | E15 | 0.2353 |
|  |  | E17 | 0.8345 |
|  | Left PICA | E13 | 0.2100 |
|  |  | E15 | 0.7841 |
|  |  | E17 | 0.5308 |
|  | Right PICA | E13 | 0.2962 |
|  |  | E15 | 0.0828 |
|  |  | E17 | 0.2962 |
| **Figure 6C**  Total artery length per age |  | E11 | 0.5402 |
|  |  | E13 | 0.6761 |
|  |  | E15 | **0.0357*** **♂ < ♀** |
|  |  | E17 | 0.2100 |
| **Figure 6C’**  Artery length per age | Left SCA | E11 | 0.5402 |
|  |  | E13 | 0.4033 |
|  |  | E15 | 0.3153 |
|  |  | E17 | 0.6761 |
|  | Right SCA | E11 | 0.3912 |
|  |  | E13 | 0.6761 |
|  |  | **E15** | **0.0081** ♂ < ♀** |
|  |  | E17 | 0.6761 |
|  | Left AICA | E13 | 0.4033 |
|  |  | E15 | 0.3153 |
|  |  | E17 | 0.6761 |
|  | Right AICA | E13 | 0.6761 |
|  |  | **E15** | **0.01371*♂ < ♀** |
|  |  | E17 | 0.0601 |
|  | Left PICA | E13 | 0.5308 |
|  |  | E15 | 0.2353 |
|  |  | E17 | 0.0601 |
|  | Right PICA | E13 | 0.2962 |
|  |  | **E15** | **0.0137* ♂ < ♀** |
|  |  | E17 | 0.8345 |
| **Figure 10C**  Network volume |  | E11 | 1.000 |
|  |  | E13 | 0.7841 |
|  |  | E15 | 0.4033 |
|  |  | **E17** | **0.0367* ♂ > ♀** |
|  |  | P0 | 0.2100 |
| **Figure 10C**  Network length |  | E11 | 1.000 |
|  |  | E13 | 0.3153 |
|  |  | E15 | 0.5308 |
|  |  | **E17** | **0.0121* ♂ > ♀** |
|  |  | P0 | 0.1436 |
| **Figure 10D**  Number of segments |  | E11 | 1.000 |
|  |  | E13 | 0.3153 |
|  |  | E15 | 0.2962 |
|  |  | **E17** | **0.0121* ♂ > ♀** |
|  |  | P0 | 0.2100 |
| **Figure10D**  Branchpoints |  | E11 | 1.000 |
|  |  | E13 | 0.6480 |
|  |  | E15 | 0.2962 |
|  |  | **E17** | **0.0121* ♂ > ♀** |
|  |  | P0 | 0.1436 |
| **Figure 10D**  Segment partitioning |  | E11 | 0.3912 |
|  |  | E13 | 0.6480 |
|  |  | E15 | 0.6761 |
|  |  | E17 | 0.0601 |
|  |  | P0 | 1.000 |
| **Figure 10E**  Mean segment volume |  | E11 | 0.5402 |
|  |  | E13 | 0.9272 |
|  |  | E15 | 0.5308 |
|  |  | E17 | 0.0946 |
|  |  | P0 | 0.5308 |
| **Figure 10E**  Mean segment length |  | E11 | 0.0661 |
|  |  | E13 | 0.9272 |
|  |  | E15 | 0.5308 |
|  |  | E17 | 0.0946 |
|  |  | P0 | 0.5308 |
| **Figure 10E**  Mean segment radius |  | E11 | 0.2703 |
|  |  | E13 | 0.6480 |
|  |  | E15 | 0.4033 |
|  |  | **E17** | **0.0367* ♂ > ♀** |
|  |  | P0 | 0.6761 |

**Supplementary table S5: Summary of the statistical analyses used for each quantitative parameter.**

| ***Analysis*** | ***Test*** | ***n*** | ***p-value*** |
| --- | --- | --- | --- |
| **Figure 6B**  Total artery volume per age | Kruskal-Wallis  Dunn’s test | E11 n=9 (5♂/4♀)  E13 n=10 (5♂/5♀)  E15 n=12 (7♂/5♀)  E17 n=10 (5♂/5♀) | Kruskal-Wallis Stat.: 15.60  p= 0.0014  E11 *vs* E13 – p= 0.0248*  E11 *vs* E15 – p= 0.0008* |
| **Figure 6B’**  Artery volume per age | Two way ANOVA Tukey post-test | E11 n=9 (5♂/4♀)  E13 n=10 (5♂/5♀)  E15 n=12 (7♂/5♀)  E17 n=10 (5♂/5♀)  Left and right arteries | **ANOVA**  Interaction F 4.614 – p<0.0001****  Row factor F 31.40 – p<0.0001****  Column factor F 3.621 – p= 0.0036**  **Tukey’s**  *Left AICA*  E13 *vs* E17 – p=0.0006***  *Right AICA*  E13 *vs* E17 – p<0.0001****  E15 *vs* E17 – p<0.0001****  *Right PICA*  E13 *vs* E17 – p=0.0175* |
| **Figure 6C**  Total artery length per age | Kruskal-Wallis  Dunn’s test | E11 n=9 (5♂/4♀)  E13 n=10 (5♂/5♀)  E15 n=12 (7♂/5♀)  E17 n=10 (5♂/5♀) | Kruskal-Wallis Stat.: 15.60  p<0,0001****  E11 *vs* E13 – p=0.0007***  E11 *vs* E15 – p=0.0037**  E13 *vs* E17 – p=0.0185 * |
| **Figure 6C’**  Artery length per age | Two way ANOVA Tukey post-test | E11 n=9 (5♂/4♀)  E13 n=10 (5♂/5♀)  E15 n=12 (7♂/5♀)  E17 n=10 (5♂/5♀)  Left and right arteries | **ANOVA**  Interaction F 9.407 – p= 0.0002***  Row factor F 18.53 – p<0.0001****  Column factor F 28.92 – p<0.0001****  **Tukey’s**  *Left SCA*  E15 *vs* E17 – p=0.0265*  *Right SCA*  E11 *vs* E17 – p=0.0239*  E13 *vs* E17 – p=0.0215*  E15 *vs* E17 – p=0.0141*  *Left AICA*  E15 *vs* E17 – p=0.0265*  *Right AICA*  E13 *vs* E17 – p= 0.0408* |
| **Figure 6D**  E11  Artery volume per individual | Wilcoxon | E11 n=9 (5♂/4♀)  Left and right arteries | p=0.6523 – ns |
| **Figure 6D**  E13  Artery volume per individual | Friedman  Dunn’s test | E13 n=10 (5♂/5♀)  Left and right arteries | **Friedman**  Friedman stat.: 13.76  p=0.0172*  **Mixed-effects analysis**  Treatment: p=0.0275*  Matching effect: p=0.0056**  V_indiv_ = 2,186 x 10^-6^  V_residual_ = 5.171 x 10^-6^  V_indiv_<V_resid_ |
| **Figure 6D**  E15  Artery volume per individual | Friedman  Dunn’s test | E15 n=12 (7♂/5♀)  Left and right arteries | **Friedman**  Friedman stat.: 22.63  p=0.0004***  **Mixed-effects analysis**  Treatment: p=0.021**  Matching effect: p=0.0069**  V_indiv_ = 3.981 x 10^-6^  V_residual_ = 1.064 x 10^-5^  V_indiv_<V_resid_ |
| **Figure 6D**  E17  Artery volume per individual | Friedman  Dunn’s test | E17 n=10 (5♂/5♀)  Left and right arteries | **Friedman**  Friedman stat.: 33.54  p<0.0001****  **Mixed-effects analysis**  Treatment : p=0.0006***  Matching effect : p=0.0671 ns  V_indiv_ = 2.682 x 10^-6^  V_residual_ = 1.172 x 10^-5^  V_indiv_<V_resid_ |
| **Figure 6D’**  E11  Artery length per individual | Wilcoxon | E11 n=9 (5♂/4♀)  Left and right arteries | p=0.5703 – ns |
| **Figure 6D’**  E13  Artery length per individual | Friedman  Dunn’s test | E13 n=10 (5♂/5♀)  Left and right arteries | **Friedman**  Friedman stat.: 39.31  p<0.0001****  Dunn  *Left AICA vs Right AICA –* p=0.0824 – ns  **Mixed-effects analysis**  Treatment: p=0.0002***  Matching effect: p<0.0001 ****  V_indiv_ = 3.130 x 10^-12^  V_residual_ = 1.307 x 10^-12^  V_indiv_>V_resid_ |
| **Figure 6D’**  E15  Artery length per individual | Friedman  Dunn’s test | E15 n=12 (7♂/5♀)  Left and right arteries | **Friedman**  Friedman stat.: 47.30  p<0.0001***  Dunn  *Left AICA vs Right AICA –* p=0.0157*  **Mixed-effects analysis**  Treatment: p<0.0001****  Matching effect: p<0.0001 ****  V_indiv_ = 6.154 x 10^-12^  V_residual_ = 1.256 x 10^-12^  V_indiv_>V_resid_ |
| **Figure 6D’**  E17  Artery length per individual | Friedman  Dunn’s test | E17 n=10 (5♂/5♀)  Left and right arteries | **Friedman**  Friedman stat.: 40.16  p<0.0001****  Dunn  *Left AICA vs Right AICA –* p=0.0824 – ns  **Mixed-effects analysis**  Treatment: p=0.0002***  Matching effect: p=0.0044**  V_indiv_ = 1.063 x 10^-13^  V_residual_ = 2.187 x 10^-13^  V_indiv_>V_resid_ |
| **Figure 7C**  Analysis of the topographic variability of the three cerebellar arterial pairs | Fisher’s exact test | E13 n=12 (7♂/5♀)  E15 n=12 (6♂/6♀)  E17 n=10 (5♂/5♀)  Left and right arteries | **Fisher exact’s test**  Sex-based morphology AICA  p=0.7249 – ns  Sex-based morphology PICA  p=0.9214 ns  Age-based morphology AICA  p=0.3576 – ns  Age-based morphology PICA  p=0.1902 – ns  Sex-based branching  p=0.4348 – ns  Age-based branching  p=0.1902  E13 *vs* E17 – p=0.05341 – ns |
| **Figure 10AB** | Principal component analysis | E11 n=9 (5♂/4♀)  E13 n=11 (6♂/5♀)  E15 n=10 (5♂/5♀)  E17 n=10 (5♂/5♀)  14 components analyzed | **Principal component analysis**  Percentage of cumulative variance  Dim1: 47.43%  Dim2: 73.92%  Dim3:89.48%  Dim4: 95.09% |
| **Figure 10C**  Network volume | Kruskal-Wallis  Dunn’s test | E11 n=9 (5♂/4♀)  E13 n=11 (6♂/5♀)  E15 n=10 (5♂/5♀)  E17 n=10 (5♂/5♀) | Kruskal-Wallis Stat.: 29.80  p<0,0001****  E11 *vs* E15 – p=0.0006***  E11 *vs* E17 – p=0.0007***  E13 *vs* E15 – p=0,0350*  E13 *vs* E17 – p=0.0427*  E15 *vs* P0 – p=0.0044**  E17 *vs* P0 – p=0.0056** |
| **Figure 10C**  Network length | Kruskal-Wallis  Dunn’s test | E11 n=9 (5♂/4♀)  E13 n=11 (6♂/5♀)  E15 n=10 (5♂/5♀)  E17 n=10 (5♂/5♀) | Kruskal-Wallis Stat.: 20.54  p=0.0004***  E11 *vs* E17 – p=0.0244*  E15 *vs* P0 – p=0.0216*  E17 *vs* P0 – p=0.0033** |
| **Figure 10D**  Number of segments | Kruskal-Wallis  Dunn’s test | E11 n=9 (5♂/4♀)  E13 n=11 (6♂/5♀)  E15 n=10 (5♂/5♀)  E17 n=10 (5♂/5♀) | Kruskal-Wallis Stat.: 18.21  P=0,0011**  E11 *vs* E17 – p=0.0105*  E13 *vs* E17 – p=0.0421*  E17 *vs* P0 – p=0.0194* |
| **Figure10D**  Branchpoints | Kruskal-Wallis  Dunn’s test | E11 n=9 (5♂/4♀)  E13 n=11 (6♂/5♀)  E15 n=10 (5♂/5♀)  E17 n=10 (5♂/5♀) | Kruskal-Wallis Stat.: 22.87  p=0,0001***  E11 *vs*. E15 – p=0.0446*  E11 *vs* E17 – p=0.0026**  E13 *vs*. E17 – p=0.0120*  E17 *vs* P0 – p=0.0059** |
| **Figure 10D**  Segment partitioning | Kruskal-Wallis  Dunn’s test | E11 n=9 (5♂/4♀)  E13 n=11 (6♂/5♀)  E15 n=10 (5♂/5♀)  E17 n=10 (5♂/5♀) | Kruskal-Wallis Stat.: 42.98  p<0,0001****  E11 *vs* E17 – p=0.0001***  E11 *vs* P0 – p<0.0001****  E13 *vs* E17 – p=0.0396*  E13 *vs* P0 – p<0.0001****  E15 *vs* P0 – p=0.0109* |
| **Figure 10E**  Mean segment volume | Kruskal-Wallis  Dunn’s test | E11 n=9 (5♂/4♀)  E13 n=11 (6♂/5♀)  E15 n=10 (5♂/5♀)  E17 n=10 (5♂/5♀) | Kruskal-Wallis Stat.: 37.21  p<0,0001****  E11 *vs* E15 – p=0.0039**  E11 *vs* E17 – p<0.0001****  E11 *vs* P0 – p<0.0001****  E13 *vs* E17 – p=0.0059**  E13 *vs* P0 – p=0.0052** |
| **Figure 10E**  Mean segment length | Kruskal-Wallis  Dunn’s test | E11 n=9 (5♂/4♀)  E13 n=11 (6♂/5♀)  E15 n=10 (5♂/5♀)  E17 n=10 (5♂/5♀) | Kruskal-Wallis Stat.: 43.13  p<0,0001****  E11 *vs* E15 – p=0.0398*  E11 *vs* E17 – p=0.0002***  E11 *vs* P0 – p<0.0001****  E13 *vs* P0 – p=0.0001***  E15 *vs* P0 – p=0.0128* |
| **Figure 10E**  Mean segment radius | Kruskal-Wallis  Dunn’s test | E11 n=9 (5♂/4♀)  E13 n=11 (6♂/5♀)  E15 n=10 (5♂/5♀)  E17 n=10 (5♂/5♀) | Kruskal-Wallis Stat.: 41.20  p<0,0001****  E11 *vs* E15 – p=0.0164*  E11 *vs* E17 – p<0.0001****  E11 *vs* P0 – p<0.0001****  E13 *vs* E17 – p=0.0135*  E13 *vs* P0 – p=0.0004*** |
